# Population-scale drug target and variant prioritization with Varformer

**DOI:** 10.64898/2026.10.16.759653

**Authors:** Aaron Wenteler, Victor Neduva, Genes & Health Research Team, Claudia P. Cabrera, Wei Wei, Michael R. Barnes

**Affiliations:** Digital Environment Research Institute, Queen Mary University of London, London, United Kingdom; William Harvey Research Institute, Queen Mary University of London, London, United Kingdom; Data, AI & Genome Sciences, Merck & Co., Inc., Cambridge, MA, United States of America; Roche Innovation Center, F. Hoffmann-La Roche Ltd, Basel, Switzerland; NIHR Barts Cardiovascular Biomedical Research Centre, Queen Mary University of London, London, United Kingdom; The Alan Turing Institute, London, United Kingdom

**Keywords:** Drug Target Prioritization, Population Genetics, Multimodal Neural Networks, Transformers

## Abstract

Human genetic variation is a powerful driver of therapeutic target prioritization, yet current models struggle to aggregate variable numbers of variants per gene and are bottlenecked by the scarcity of validated targets. We present Varformer, a multimodal transformer that uses cross-modal attention to integrate contextual representations of missense variants with gene-level features. Trained via positive-unlabeled learning on clinically validated targets across four genetic ancestry groups, Varformer achieves a precision@20 of 0.95, 0.86, and 0.48 across three benchmarks of increasing difficulty. It surpasses a logistic regression ablation by 20% and DrugnomeAI by 23% on the hardest benchmark. Cross-population rank normalization isolates targets with consistent genetic support across ancestry groups. The variant encoder also scores individual variants. Across 1,164 carrier–variant pairs in Genes & Health, Varformer ranked the known pathogenic variant in the carrier’s top 20 in 57% of pairs. Varformer is available as an open-source framework for unified target and variant prioritization.

## Main

Identifying safe and efficacious therapeutic targets remains one of the most consequential challenges in drug discovery and accounts for a substantial proportion of clinical trial failures [1, 2]. Human genetics is an increasingly important source of evidence for target discovery, yet allele frequencies and genetic composition differ across ancestrally diverse populations and influence both drug efficacy and safety [3–7]. Prioritization frameworks that ignore population-specific genetic architecture risk perpetuating therapeutic inequities.

Missense variation is a major driver of human disease [8, 9], and deep-learning models now predict missense pathogenicity at high resolution [10–12]. Existing target prioritization frameworks nevertheless collapse this information into handcrafted gene-level summary statistics, which limits their ability to capture the heterogeneous functional landscape of naturally occurring variation. A central methodological challenge is therefore to learn directly from variable-length sets of missense variants while integrating complementary gene-level biological information.

Target prioritization is naturally a positive-unlabeled learning problem. Few successful targets are known, most candidate genes are unlabeled, and clinical trial failure does not necessarily indicate an ineffective target, because outcomes also depend on trial design, endpoint selection and commercial considerations. We therefore train Varformer with a non-negative positive-unlabeled (nnPU) learning objective, which estimates classification risk from positive and unlabeled examples without explicit negative labels [13–15].

Existing machine learning methods for target discovery differ in the label that they train against. Some score human genetic evidence for disease association [16–19], which is necessary but not sufficient for therapeutic value. Others predict the outcome of a clinical trial for a target that has already been chosen [20, 21], or learn general-purpose representations of genomic sequence, cellular state or clinical data whose pre-training objective is not tied to therapeutic potential [22–24]. DrugnomeAI [25], built on the Mantis-ML framework [26, 27], supervises directly on clinical precedent and is, to our knowledge, the only publicly available disease-agnostic, gene-level target prioritization model trained on clinical trial approvals. It is therefore closest to the work presented here, but its precomputed gene-level features preclude predictions tailored to populations of different genetic backgrounds. Across these approaches, the genetic evidence comes predominantly from cohorts of European ancestry and none learns from population-scale variation directly, so no existing method can prioritize targets differentially across ancestries.

Here we present Varformer, a multimodal transformer that represents population-scale missense variation at both the gene and variant levels (Fig. 1). A shared variant encoder learns from the alleles observed in each gene. Cross-attention integrates these representations with gene-level biological features to predict therapeutic target potential, while a token-level readout assigns a prioritization score to each retained allele. We evaluate gene predictions in four ancestrally distinct cohorts—African (AFR), Admixed American (AMR), Non-Finnish European (NFE) and South Asian (SAS)— and use cross-population scores to identify targets with consistent genetic support. In the Genes & Health (G&H) SAS cohort, we evaluate the variant output for retrieving known pathogenic alleles from individual candidate lists and for identifying variants associated with measured traits.

**Fig. 1:**
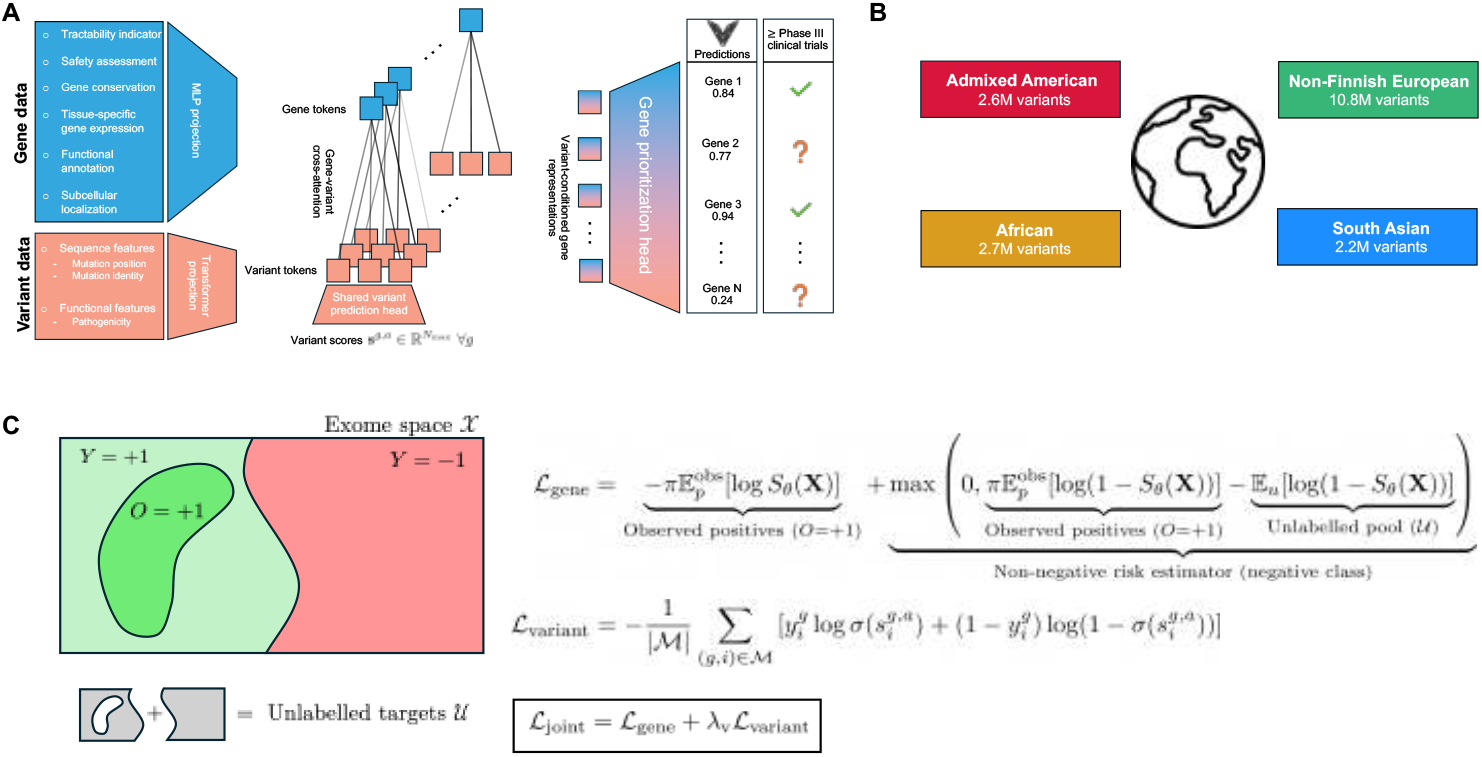
Varformer integrates gene features and population-scale missense variation for gene and variant prioritization. **A)** Varformer’s architecture. Gene features are projected into gene representations, while a Transformer encoder produces contextual variant tokens. Within each gene, cross-attention uses the gene representation to aggregate information from its variant tokens to the gene-level. The gene prioritization head takes these gene-level representations, and produces *S*_*θ*_(**x**^*g,a*^) ∈ [0, 1] for gene *g* in population *a*. A shared linear variant head prioritizes the contextual variant tokens before cross-attention and produces the variant score vector 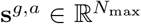, with *N*_max_ = 1,024 and padding positions masked. The variant-head is a single linear layer, so parameters are shared between every token and gene for one population model. **B)** Populations used to train the gene-prioritization models: Admixed American (AMR, red), South Asian (SAS, blue), Non-Finnish European (NFE, green), and African/African-American (AFR, gold). Boxes indicate the corresponding variant counts. **C)** Gene and variant supervision. The schematic illustrates the input space *X*, with latent positive and negative genes (*Y* ∈ {−1, +1}), observed positive genes (*O* = +1), i.e. drug targets that reached at least Phase III of clinical trials, and an unlabeled pool *U* containing both unobserved positives and negatives. The gene loss combines observed-positive and unlabeled examples and clips the estimated negative-class risk at zero for stability. The variant loss is the binary cross-entropy over eligible tokens ℳ, using allele frequency-derived labels 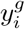 and sigmoid-transformed token logits. Joint training minimizes ℒ_joint_ = ℒ_gene_ + *λ*_*v*_ℒ_variant_.

## Results

Varformer combines gene-level features, including tractability, safety and tissue-specific expression, with population-specific missense variation. Its population variant characterization (PVC) module produces a contextual embedding for each retained variant. These embeddings feed a gene-level prioritization score through cross-attention and a variant-level score through a shared prediction head. The gene objective uses nnPU learning on clinical trial status, and the variant objective uses allele frequency as a proxy for selective constraint; both update the variant encoder jointly (Methods, Model training and Evaluation of variant prioritization).

### Varformer learns representations of therapeutic potential

To examine how Varformer organizes genes by therapeutic potential, we compared UMAP projections of gene-level variant summaries with the learned embeddings in each population (Fig. 2A–D). In the input summaries, known therapeutic targets were dispersed among unlabeled genes. In the learned embeddings, they concentrated in distinct regions of the latent space in all populations.

**Fig. 2:**
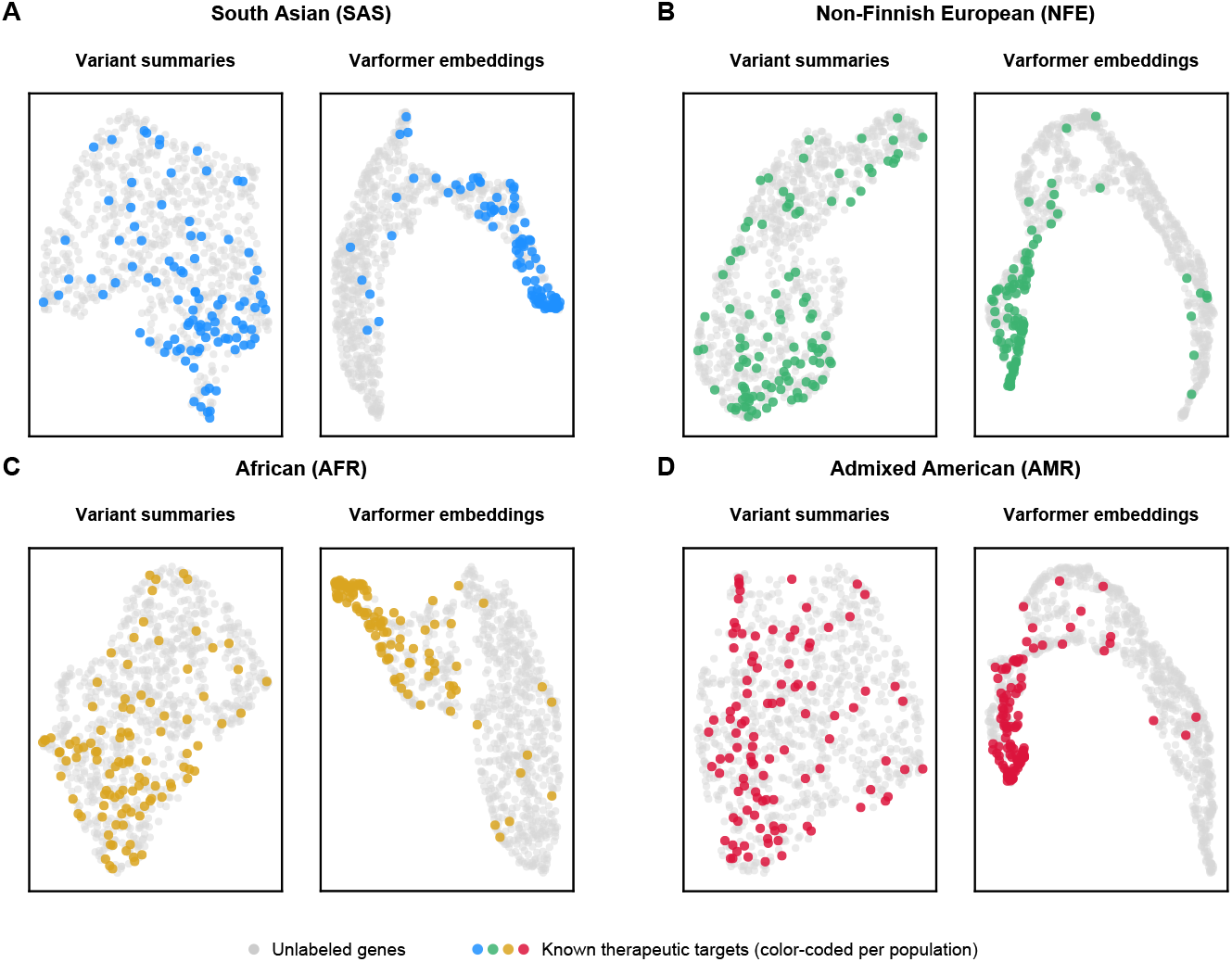
Varformer embeddings concentrate approved drug targets in every cohort. UMAP projections for **A)** South Asian, **B)** Non-Finnish European, **C)** African and **D)** Admixed American populations. Each panel compares gene-level variant summary statistics (left) with Varformer embeddings (right), colored by drug target approval status. Approved drug targets are shown in the population color used in the other figures (SAS blue, NFE green, AFR gold, AMR red) and show tighter and more delineated clustering in Varformer embedding space compared to the input summaries. Unlabeled genes are shown in gray.

We quantified this organization by partitioning the standardized 256-dimensional embeddings into two clusters within each population (Supplementary Fig. 1A; Supplementary Note 1). One cluster contained a higher proportion of known drug targets: 38.8% in SAS, 39.4% in NFE, 42.1% in AFR and 40.5% in AMR, compared with 2.2%, 2.2%, 2.5% and 2.1%, respectively, in the other cluster (Supplementary Fig. 1C). This corresponds to an approximately 17- to 19-fold enrichment in target density. Cluster membership showed no significant association with benchmark identity (*χ*^2^ test, *p >* 0.05; Supplementary Fig. 1B). Among unlabeled genes, the target-enriched cluster included genes with three or four chemical-tractability annotations, whereas 89–92% of unlabeled genes in the other cluster had none (Supplementary Fig. 1D). These annotations are among the gene inputs, so their distribution shows how chemical-tractability information is reflected in Varformer’s representation space.

### Varformer prioritizes drug targets across populations

We evaluated Varformer on three progressively challenging benchmarks: *Pfam Druggable Targets* (Pfam), *Recently Approved Targets* and *Chemically Putative Targets* (Pharos). We compared it with a random baseline, a logistic regression model on hand-crafted summary features (Varformer-LR), which isolates the value of the architecture, and DrugnomeAI. The default feature set of DrugnomeAI includes drug–gene interaction records from DGIdb and the Comparative Toxicogenomics Database (CTD), which exist for a gene largely because that gene is already a drug target and therefore reveal the label that the benchmarks test. To prevent this leakage, we trained DrugnomeAI on the positive labels of Varformer without the DGIdb feature and without the CTD features that record chemical–gene interactions (Methods, Benchmarking setup and evaluation strategy; Supplementary Note 9). We assessed the top of the ranked list with precision@20, ranking quality over each complete benchmark with the Spearman correlation coefficient (*ρ*), and classification quality with the F1 score. Values are reported as mean ± standard deviation across five seeds in each of the four populations (Supplementary Table 1).

At the top of the ranked list, Varformer’s precision@20 was 0.95 ± 0.04 on Pfam, 0.86±0.05 on Recently Approved and 0.48±0.08 on Pharos, compared with 0.95±0.05, 0.80 ± 0.05 and 0.40 ± 0.07 for Varformer-LR and 0.91 ± 0.03, 0.84 ± 0.04 and 0.39 ± 0.06 for DrugnomeAI (Fig. 3A–C). Varformer’s precision@20 was equal to that of Varformer-LR on Pfam and 6.9% and 20.0% higher on Recently Approved and Pharos, and it was 3.8%, 1.8% and 23.1% higher than that of DrugnomeAI on the three benchmarks, respectively. The advantage was largest on Pharos, the most difficult benchmark, where Varformer exceeded both Varformer-LR and DrugnomeAI (adjusted *q* = 0.005 for each) and recovered approximately two additional benchmark positives per 20-gene shortlist. With a positive fraction of approximately 0.11 in this benchmark, its precision@20 represents a roughly fourfold enrichment over random ranking. Ranking quality over the complete benchmark sets showed a more mixed pattern (Fig. 3D; Supplementary Fig. 2). On Pfam, the Spearman correlation of Varformer did not differ from that of DrugnomeAI or Varformer-LR (0.53 ± 0.01 for each model). On Recently Approved, DrugnomeAI had the highest Spearman correlation (0.53 ± 0.01, compared with 0.50 ± 0.01 for Varformer-LR and 0.48 ± 0.01 for Varformer). On Pharos, Varformer (0.39 ± 0.02) exceeded both DrugnomeAI (0.24 ± 0.01) and Varformer-LR (0.37 ± 0.01; *q <* 0.001 for each). F1 scores were higher for Varformer than for Varformer-LR on all three benchmarks (*q* ≤ 0.004). Compared with DrugnomeAI, Varformer had a higher F1 score on Pharos (0.44 ± 0.05 compared with 0.33 ± 0.04; *q <* 0.001), and the F1 comparisons on Pfam (0.87 ± 0.04 compared with 0.85 ± 0.03) and on Recently Approved (0.75 ± 0.04 compared with 0.74 ± 0.04) were inconclusive. Across all four populations, Varformer’s predictions concentrate near the extremes of the [0, 1] interval, with most of the mass at low scores, consistent with the low prior probability of clinical success (Fig. 3E).

**Fig. 3:**
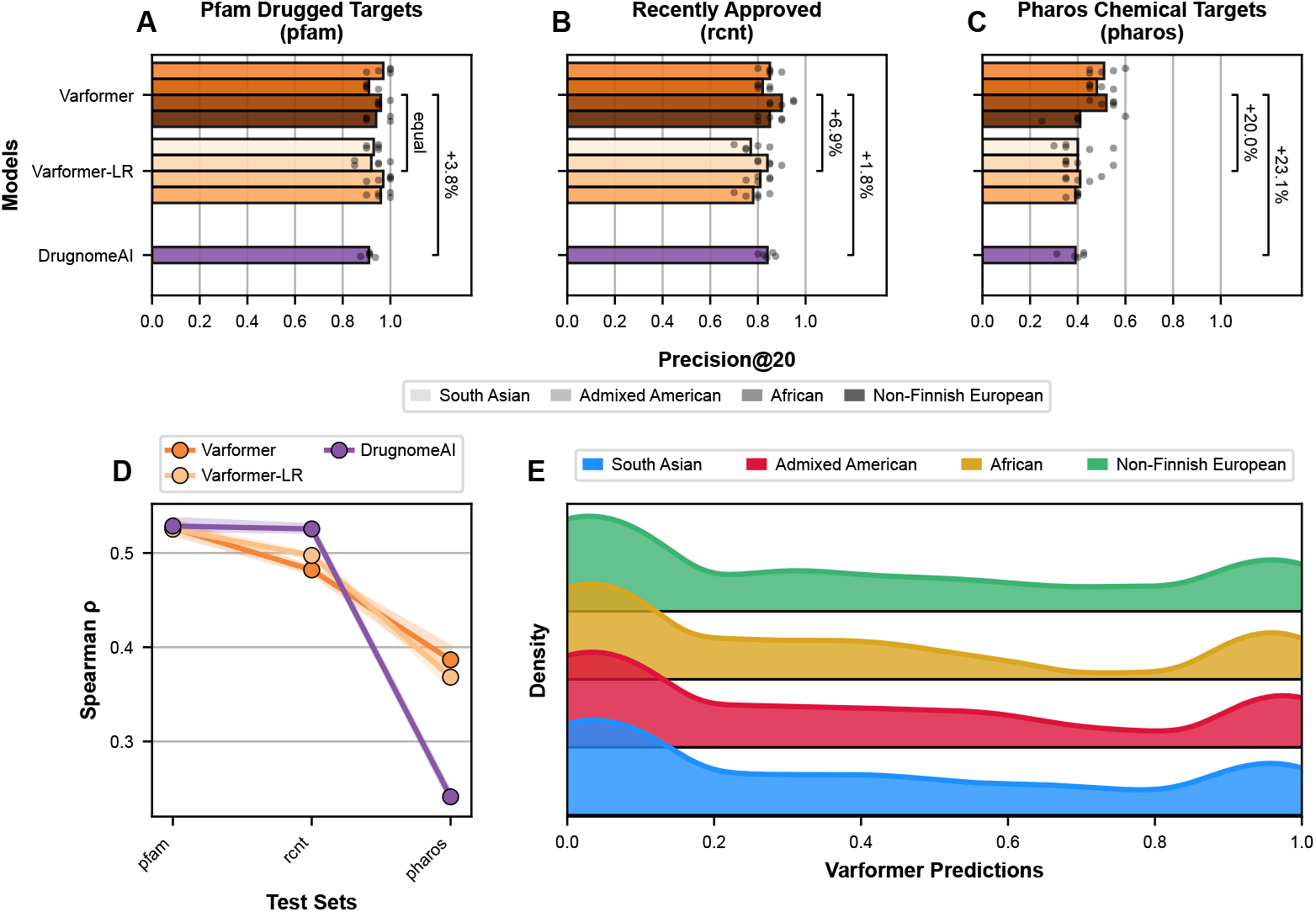
**A-C)** Precision@20 on the three benchmarks for Varformer, a logistic regression variant of Varformer and DrugnomeAI. Bars show the mean across five seeds and points show individual seeds. Varformer and Varformer-LR are shown for each population; for DrugnomeAI, trained without drug–gene interaction features, each point is the mean of one seed across the four population-specific benchmark sets. **D)** Spearman correlation between predicted scores and benchmark labels across the three test sets for all benchmarked models. Lines show the mean across 20 runs and bands show bootstrap 95% confidence intervals. Color coding is consistent with A-C. **E)** Density distribution of Varformer prediction scores for the benchmark genes in each population. The density axis is logarithmic and spans two orders of magnitude below the peak of each distribution.

### De-risking drug targets against population-level genetic variation

To identify targets whose prioritization is robust to inter-population genetic variation, we converted Varformer scores into rank-normalized scores. These place each target on a [0, 1] quantile scale within its own population and are therefore comparable across populations. Among unlabeled genes scored in all four populations, the ten candidates with the highest mean cross-population percentile range from 98.96 to 99.66, with across-population standard deviations of 0.06 to 0.72 percentile points, and every candidate remains at or above the 98.28th percentile in its lowest-ranking population (Fig. 4A).

**Fig. 4:**
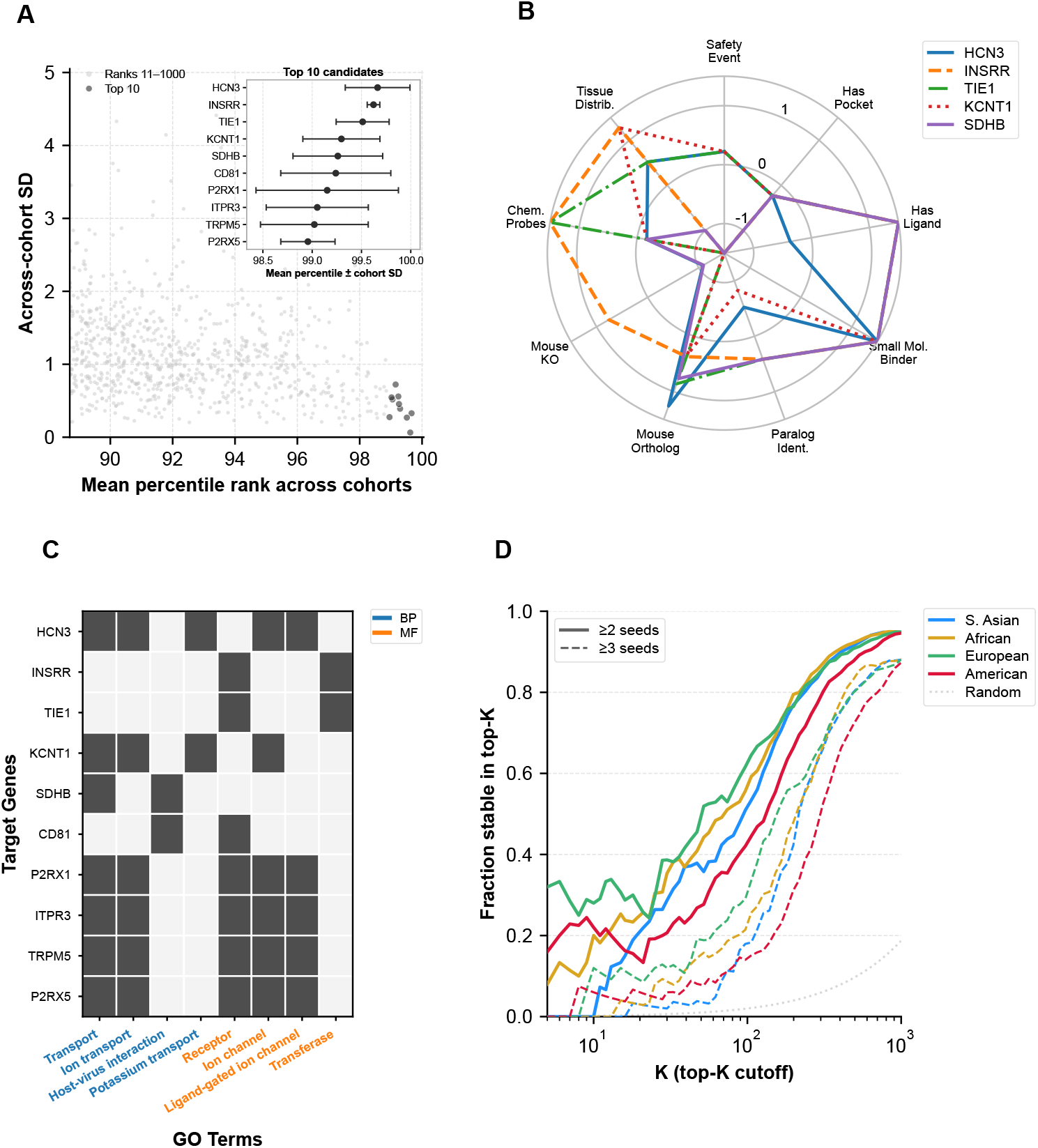
De-risking and characterization of cross-population drug-target predictions. **A)** The 1,000 genes with the highest mean percentile rank across populations, plotted against the SD of their population-specific percentiles. The ten candidates with the highest mean percentile are shown in black and highlighted in the inset. **B)** Radar plot of the nine Open Targets gene characterization features used as model inputs, for the top five candidates. **C)** Binary GO membership heatmap for the top ten candidates. Term-label colors distinguish biological process (BP; blue) from molecular function (MF; orange); filled cells indicate membership of the corresponding GO term. **D)** Top-*K* stability across five training seeds. Curves show the mean fraction of each seed’s top-*K* set belonging to genes that recur in at least two seeds (solid) or at least three seeds (dashed), with a random ranking baseline (dotted gray).

The five leading candidates—*HCN3, INSRR, TIE1, KCNT1* and *SDHB* —share small-molecule-binder annotations, and four of them have ligand annotations, but they differ in safety-event annotations, orthology, tissue distribution and chemical-probe evidence (Fig. 4B; Supplementary Note 2). Similarly high cross-population rankings therefore encompass distinct combinations of features relevant to target development. The Gene Ontology annotations of the top ten candidates show a shared functional theme (Fig. 4C): transport and receptor activity each annotate seven candidates, and ion transport and ion-channel activity six each. Ion channels and several receptor classes are well-established drug target families [28, 29].

We assessed ranking reproducibility with a top-*K* stability analysis across five independent training seeds (Fig. 4D). At *K* = 1,000, 94.6–94.9% of genes in a seed’s top-*K* set recur in at least two seeds, and 87.3–88.3% recur in at least three. The cross-population and seed-recurrence analyses thus provide complementary measures of robustness.

### Prospective validation of Varformer drug target predictions

To assess prospective validity, we performed a time-stratified evaluation with a training cutoff of 29 June 2023, corresponding to Open Targets release 23.06, and an outcome snapshot from release 26.06. Of the 332 genes recorded at Phase I/II at the training cutoff, 243 belonged to the training-unlabeled pool and had ensemble predictions in at least two populations. We summarized each gene by its mean rank-normalized score and applied the prior-derived threshold for the number of populations in which the gene was scored (Methods, Time-stratified prospective validation). Sixteen targets had major progression events between June 2023 and June 2026: 13 moved from Phase I/II to Phase III or approval, and 3 failed owing to Phase II efficacy or safety concerns. Targets without a resolved outcome were not treated as negatives. Ten of the 13 progression events were above the respective threshold, but only one of the three failures was below it, giving concordant predictions for 11 of 16 genes (68.8%).

For example, *KDM1A* and *TMPRSS6* were above the four-population threshold before programs engaging these targets advanced to Phase III. *TREM2* and *OSMR* were also above this threshold, although Phase II trials of agents against them subsequently failed, so the gene-level score did not anticipate these program-specific failures (Supplementary Note 3).

### Varformer prioritizes missense variants for individual-level follow-up

The variant representations underlying Varformer’s gene predictions provide a score for each retained missense token (Fig. 1A). Averaging across five SAS model seeds and repeated gene-associated annotations gives one variant score, 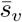, per DNA variant. We evaluated these scores in G&H, a cohort of British Bangladeshi and Pakistani participants [30]. Among 2,477 pathogenic or likely pathogenic and 15,580 benign or likely benign rare variants (cohort allele frequency, AF *<* 10^−3^), 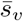 achieved an AUROC of 0.890 and average precision of 0.627, compared with 0.905 and 0.644 for AlphaMissense and 0.606 and 0.177 for Varformer’s gene score, which assigns an equal score to every variant in the same gene (Fig. 5A). Across all frequencies (2,521 pathogenic or likely pathogenic and 25,777 benign or likely benign variants), 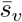 achieved an AUROC of 0.891 and average precision of 0.544, compared with 0.902 and 0.551 for AlphaMissense. REVEL achieved the highest average precision among individual scores (0.647; Fig. 5B).

**Fig. 5:**
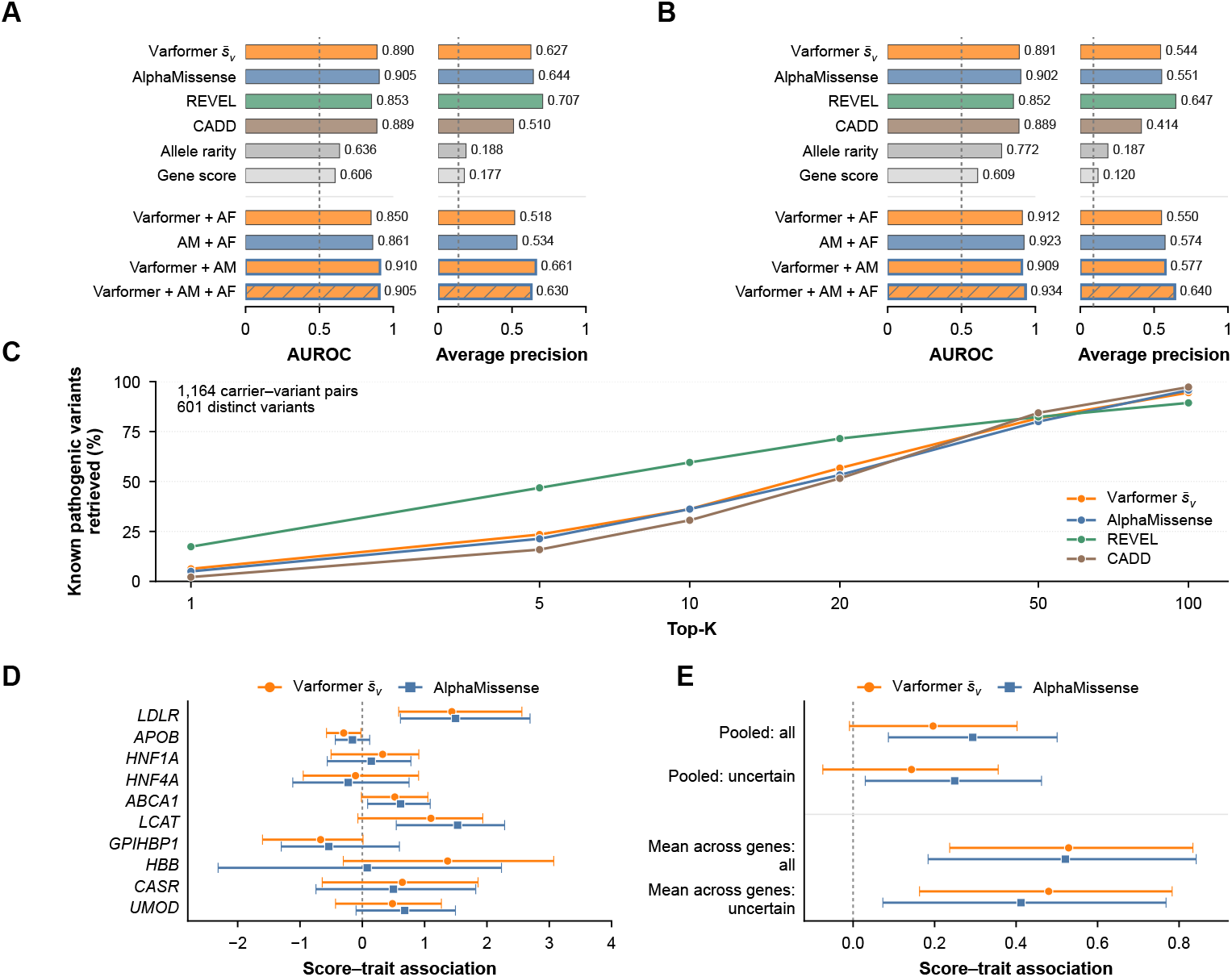
Variant prioritization and phenotype associations of Varformer’s variant scores. **A)** AUROC and average precision (AP) among 2,477 pathogenic/-likely pathogenic and 15,580 benign/likely benign rare variants (cohort AF *<* 10^−3^). Dashed references: AUROC 0.5 and AP 0.137. Allele rarity is the negated cohort AF; the Varformer gene score assigns one value to all variants in a gene. **B)** All-frequency classification (2,521 pathogenic/likely pathogenic; 25,777 benign/likely benign variants), evaluated as in A. Both panels include equal-weight percentile combinations. AM denotes AlphaMissense; AF denotes allele rarity. Blue outlines denote Varformer– AM combinations; gray diagonal hatching denotes the three-way score. The dashed AP reference is 0.089. A–C share ClinVar evidence. **C)** Known pathogenic variants (ClinVar pathogenic/likely pathogenic) retrieved at six top-*K* cutoffs across 1,164 carrier–variant pairs and 601 distinct variants. **D)** Gene-specific 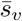 and AlphaMissense score–trait associations in the uncertain subset, with pointwise 95% intervals from 50,000 variant-bootstrap draws. Adjusted intervals for 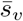 and paired differences are reported in Supplementary Table 8. **E)** Pooled score–trait associations and mean associations across ten eligible genes 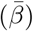. All: *n* = 1, 801, 723 variants; uncertain: *n* = 1, 608, 675 variants. “Uncertain” includes unannotated, uncertain or conflicting classifications. Lines show pointwise 95% variant-bootstrap intervals conditional on the analyzed panel. In D and E, score–trait associations measure adjusted, direction-oriented trait change across the 0–1 within-gene score-percentile scale; positive values indicate greater trait deviations for higher-ranked variants.

To assess whether the methods provide complementary information, we combined their percentile ranks with equal weights. Combining 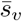 with AlphaMissense improved classification over either score alone in both variant sets: the combination achieved an AUROC of 0.910 and average precision of 0.661 among rare variants, and 0.909 and 0.577 across all frequencies (Supplementary Table 7). Adding allele rarity, which we define as the negated cohort allele frequency so that rarer variants receive higher scores, gave the highest AUROC across all frequencies (0.934) but reduced both metrics among rare variants. The three-way score also exceeded a control that assigned two-thirds weight to AlphaMissense and one-third to rarity in both sets, which supports a contribution from Varformer beyond a reweighting of allele frequency.

Across 1,164 carrier–variant pairs representing 601 distinct ClinVar pathogenic or likely pathogenic variants, 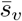 retrieved the known pathogenic variant within the carrier’s top 20 candidates in 56.7% of pairs, compared with 53.3% for AlphaMissense; both scores achieved 36.2% recall at the top-10 cutoff (Fig. 5C). REVEL achieved higher recall at both cutoffs (59.5% at *k* = 10 and 71.5% at *k* = 20). Candidate lists contained a median of 143 scoreable rare missense variants. Varformer covered 66.8% of carried rare variants and 72.9% of known pathogenic variants, because its encoder retains at most 1,024 variant tokens per gene.

We also examined whether 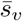 captures phenotype-relevant variation beyond confidently classified ClinVar alleles. We related within-gene score ranks to covariate-adjusted traits in carriers of rare missense variants across 15 gene–trait pairs, of which ten genes had sufficient carriers and distinct variants for gene-specific estimation (Fig. 5D).

In an exploratory summary that gives each eligible gene equal weight, the mean score– trait association across genes was 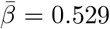 for all qualifying variants and 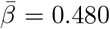 for unannotated, uncertain or conflicting variants, compared with 0.521 and 0.412 for AlphaMissense (Fig. 5E). The carrier-weighted pooled association was weaker (0.143 among the 1,608 observations carrying variants without confident classifications), and no gene showed a statistically resolved advantage over AlphaMissense. The association is therefore not uniform across genes (Supplementary Notes 4–5).

## Discussion

Varformer shifts drug target prioritization from gene-level aggregates and GWAS-led signals to individual variants modeled directly at population scale. By combining a multimodal transformer with positive-unlabeled learning, we show that clinical trial success likelihood can be estimated from genetic variation, multi-omic descriptors and functional annotations.

Within each population, Varformer’s embeddings distinguish approved targets from the unlabeled exome (Fig. 2), and a cluster with a 17- to 19-fold enrichment for known targets recurs in all four ancestries. This organization suggests that Varformer captures functional relationships relevant to therapeutic potential. Handcrafted statistics of mutational burden and spatial variant clustering are highly informative: Varformer-LR approached Varformer on the two easier benchmarks. The transformer architecture adds value most clearly on the most difficult benchmark, where Varformer exceeded Varformer-LR in precision@20, Spearman correlation and F1 score (Fig. 3). A plausible source of this gain is cross-modal attention, which conditions variant representations on a gene’s safety and druggability profile.

The nnPU objective addresses the selection bias inherent to target prioritization. By treating the unlabeled pool as a mixture of viable-but-untested targets rather than definitive negatives, Varformer surfaces candidates that lack clinical trial annotation for reasons unrelated to their biological validity. Our cross-population analysis identifies targets whose success proxy is both high and stable across ancestrally diverse populations (Fig. 4). This robustness provides computational evidence that these candidates are de-risked against population-level genetic variation, in the specific sense that their prioritization does not depend on one cohort’s variation profile. It does not establish population-independent efficacy or safety, which require experimental and clinical validation.

In the prospective evaluation, 10 of 13 targets that progressed to Phase III or approval were above their threshold, but so were two of the three targets that failed in Phase II. *KDM1A* carries few missense variants and would be undervalued by variant burden alone, but its gene-level druggability and functional features supported a high cross-population rank before its progression to Phase III. Programs targeting *TLR8* and *WEE1* advanced to Phase III despite separate futility- or safety-related terminations, and an *ITGB6* -directed program missed its primary efficacy endpoint after entering Phase III. Varformer’s cross-population gene score therefore reflects whether a gene is a plausible target, and not the molecule-, indication-, economic- and trial-specific factors that also determine the outcome of a clinical trial.

Varformer’s variant output makes the learned representation useful at the level of individual alleles. The variant tokens can be used to rank the missense variants carried by a participant or to examine the highest-scoring observed variants within a gene of interest. Our individual-level analyses in G&H focused on rare variants (cohort AF *<* 10^−3^), which represent a subset of variants where frequency alone provides little discrimination between pathogenic and benign. 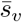 concentrated known pathogenic alleles among its top predictions and was associated with measured trait variation when eligible genes were weighted equally. The *LDLR* result links higher scores to higher LDL cholesterol among carriers whose variants lack confident clinical classifications. Although AlphaMissense is an input to Varformer’s variant encoder, combining it with 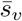 improved classification over either score alone, which indicates that the learned representation adds complementary information.

These analyses also have limits. REVEL retrieved known pathogenic variants more often than 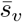 or AlphaMissense. REVEL is trained on curated disease variants and is used as evidence in clinical variant classification [31, 32], so its advantage on ClinVar-defined targets may partly reflect overlap with these labels, whereas Varformer’s variant output uses no clinical labels. Furthermore, because the variant score is learned without clinical labels and has not been calibrated against clinical classifications, it should only be interpreted as a prioritization score and not as calibrated evidence for clinical diagnosis. The carrier-weighted pooled phenotype association was uncertain, in part because *APOB*, in which variants can have opposite effects on the trait, contributed 45% of these carrier observations (721 of 1,608; Supplementary Note 5). The phenotype comparisons did not establish superiority over AlphaMissense, and the evidence instead supports combining the AlphaMissense score and 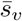. Coverage is limited to retained population-observed missense variants, so expanding the scoring subset and evaluating shortlisted variants experimentally are important next steps.

Because Varformer is disease-agnostic, a single model can prioritize across the entire exome without restriction to one indication, and the cross-population de-risking metric and the variant-level prioritization operate uniformly across disease biology. The trade-off is that the model does not account for hurdles specific to a therapeutic area: targets in central nervous system disorders, for example, face blood–brain-barrier permeability and neurobiological complexity that raise failure rates independently of genetic validity. The present model also considers missense coding variation only. Future work could add regulatory variation and disease context, for example through disease ontologies or tissue-specific regulatory networks.

By linking the gene and the variant in one model, Varformer turns sequencing data from diverse populations into a single basis for deciding which targets and which variants to pursue for therapeutic development. More broadly, the core formulation of learning to rank genes and variants from population-scale genomic features under sparse positive labels is not specific to drug target prioritization. The same approach could prioritize candidate causal genes at GWAS or rare-variant disease loci, distinguish cancer driver genes from passengers, and identify gene biomarkers of disease or treatment response. As sequencing efforts grow in scale and diversity, the amount of variation available to such models will increase, and with it their capacity to guide therapeutic discovery.

## Methods

### Data

#### Multi-ancestry population-scale exome datasets

To capture the impact of genetic variation across diverse ancestries, we utilized population-scale exome data from four major continental groups: South Asian (SAS), Non-Finnish European (NFE), Admixed American (AMR), and African/African-American (AFR). While genetic research has historically been biased toward populations of European descent, it is well-established that drug response and disease risk vary substantially across ethnic groups [33–36]. Incorporating such diversity is important for developing treatments that are both precise and more equitable.

The South Asian (SAS) cohort is derived from the Genes & Health (G&H) dataset [30], consisting of over 44,000 individuals of British Pakistani and Bangladeshi heritage. This population exhibits a higher burden of complex diseases, such as coronary heart disease and type 2 diabetes [37, 38], and is characterized by elevated levels of autozygosity [39]. These genetic characteristics offer a means of identifying population-specific therapeutic targets.

Variant data for the Non-Finnish European (NFE), Admixed American (AMR), and African/African-American (AFR) populations were obtained from the Genome Aggregation Database (gnomAD v4.1) [40]. gnomAD aggregates data from numerous large-scale sequencing projects to provide a standardized reference of human genetic variation. Across all populations, our analysis restricted focus to missense variants, which represent the most common class of functionally impactful coding variation. In total, over 2 million potential variant consequences were processed for the SAS cohort (Supplementary Fig. 3), alongside millions of variants from the gnomAD populations, ensuring that Varformer’s prioritization signals are evaluated across a broad spectrum of human genetic diversity.

#### Holdout test datasets

We test Varformer on three curated test datasets, each covering different dimensions of difficulty associated with predicting clinical trial success given representations of drug targets.

The first independent test set, termed *Pfam Druggable Targets (Pfam)*, was designed to evaluate the model’s performance on a well-defined collection of protein targets. This set includes 28 proteins from four historically druggable families: 8 G-Protein Coupled Receptors (GPCRs), 6 ion channels, 6 nuclear receptors, and 8 kinases. These classifications are informed by the Pfam database [41], a widely used resource for organizing protein sequences into families and domains.

The second test set we refer to as *Recently Approved Targets (Recent)*. These are drug targets of recently launched drugs, with launch years from 2020 to 2024. The rationale is that as the well-known druggable families have gotten depleted over the years, drug discovery efforts have shifted toward less conventional targets. Consequently, evaluating the model’s performance on recently approved drug targets allows us to assess its ability to identify a more diverse and less conventional set of targets. We classify this test set as one step up in difficulty compared to the well-known druggable protein families, as these targets are still proven to be druggable via FDA approval, but less pronounced than the druggable protein families. These targets are obtained from Citeline’s Pharmaprojects database by selecting the 32 most recently launched targets as of June 2024, of which 30 are present in our gene feature data.

The final and most challenging test set is the *Chemically Putative Targets (Pharos)* set. This group includes putative drug targets associated with highly potent small-molecule ligands that have not yet achieved FDA approval. The source data originates from Pharos v3.18.1, where these targets are classified as Tchem [42]. While the dataset initially contained approximately 2,000 chemical targets, we reduced this number to 50 to maintain consistency with the two other datasets and to avoid withholding excessive information from the training data. Additional filtering was performed to exclude targets unresolved at the gene level, resulting in a final set of 46 test targets. Because none of these targets have successfully passed clinical trials, their relevance remains less established. Consequently, this test set provides the most stringent assessment of the model’s predictive ability. Successfully identifying these putative chemical targets would demonstrate the model’s capacity to uncover promising, yet unapproved drug targets.

The benchmark negatives are unlabeled genes that are unlikely to be favorable drug targets. We selected the most genetically constrained genes (Open Targets genetic constraint *<* −0.90), the most loss-of-function-intolerant genes in the genome, and excluded all benchmark positives. For each population and seed, we sampled the benchmark negatives at random from the unlabeled genes that meet this criterion (722 negatives in SAS, 830 in AFR and in AMR, and 833 in NFE) and divided them between the three benchmark sets in proportion to the number of positives in each set, giving a positive fraction of approximately 0.11 to 0.12. The assumption that highly constrained genes are poor drug targets relies on the observation that such genes are often essential and broadly active across healthy human tissue, leading to an increased risk of off-target toxicity [43, 44]. However, constraint or essentiality in and of itself does not necessarily exclude a target from being successfully druggable [45].

A comprehensive list of the targets and their respective sources is provided in Supplementary Table 3.

### Data preprocessing and feature generation

To capture the biological factors underlying drug safety and efficacy, Varformer integrates data across two primary modules: Gene Characterization (GC) and Population Variant Characterization (PVC). This multimodal architecture is designed to represent a target’s properties across genomic, functional, and population-genetic scales. These modules are learned simultaneously and ensembled in Varformer’s output layer to obtain the final gene-level predictions. A complete description of all input features and their data sources is provided in Supplementary Table 2

#### Gene characterization module feature generation (GC)

The Gene Characterization (GC) module integrates global, gene-level features to characterize potential therapeutic targets across molecular, functional, and spatial domains. This module comprises two distinct feature sets that are processed in parallel and fused into a shared gene representation.

##### Target tractability and safety features

This set captures nine feature types designed to evaluate a target’s feasibility and safety profile. The features are sourced from the Open Targets Target Prioritization dataset [17]. Tractability is described by four binary indicators. `hasLigand` indicates at least one high-quality ligand in ChEMBL according to the Open Targets tractability assessment, `hasSmallMoleculeBinder` indicates a structure of the target co-crystallized with a small molecule in the Protein Data Bank, `hasPocket` indicates a DrugE-BIlity score of at least 0.7, which predicts a high-quality binding pocket, and `hasHighQualityChemicalProbes` indicates at least one high-quality chemical probe in the Probes & Drugs portal. These indicators describe chemical matter and protein structure. The Open Targets clinical precedence assessment, which records whether a gene is targeted by a drug at any clinical stage, is not a model input and is used only to define the training labels (Training labels). Target safety is reflected in `hasSafetyEvent`, which captures known safety liabilities. Functional assessment is captured through the `mouseKOScore`, while the gene’s degree of conservation is encoded by `paralogMaxIdentityPercentage` and `mouseOrthologMaxIdentityPercentage`. Finally, `tissueDistribution` measures the extent of baseline expression across human tissues. The Open Targets `geneticConstraint` feature is excluded from the model inputs, because it defines the benchmark negatives (Holdout test datasets). Altogether, these nine features are designed to provide an interpretable view of gene-level properties relevant to drug development success, spanning molecular, genomic, and functional domains. See Supplementary Note 7 for details on feature normalization and feature value interpretation.

##### Target function and expression features

This set integrates 527 binary features (524 in the SAS cohort, in which three biological process terms annotate no gene) that characterize the physiological role and spatial distribution of potential drug targets. Functional features were derived from the Gene Ontology (GO) biological process and molecular function categories sourced from UniProt [46], providing a high-level representation of a gene product’s molecular activity. These were complemented by subcellular localization data from the Human Protein Atlas (HPA) [47], where localization to the plasma membrane or extracellular space serves as a proxy for therapeutic accessibility. To capture the physiological context of target activity, we integrated tissue-specific abundance data from both HPA (immunohistochemistry) and the Genotype-Tissue Expression (GTEx) project (RNA-seq). Each of these features was transformed into a binary representation, where a value of 1 indicates evidence of a target’s association with a specific ontology term, subcellular compartment, or tissue type, and 0 indicates absence.

#### Population variant characterization module feature generation (PVC)

The PVC module encodes ancestral missense variation into contextual variant representations. These representations support both gene-level prioritization and direct variant scoring. Specifically, for each gene *g*, the PVC module encodes ancestral missense variation as a sequence of variant-level feature vectors. Each variant token, indexed by *i*, is represented by three primary features: (i) the AlphaMissense pathogenicity score *α*_*i*_ ∈ [0, 1], (ii) a model-position index 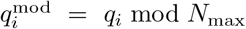 obtained from the reported amino acid coordinate *q*_*i*_, and (iii) a unique integer index *m*_*i*_ representing the specific amino acid substitution (REF → ALT). Here, *N*_max_ = 1, 024, so the coordinates are mapped into the finite positional-encoding index range 0–1,023.

To accommodate the varying number of variants per gene across different ancestral cohorts while maintaining a fixed-size input for the transformer architecture, we employ a pathogenicity-informed selection strategy. Variants within each gene are sorted in descending order of their AlphaMissense pathogenicity scores, and the top *N*_max_ = 1, 024 most pathogenic variants are retained. Genes with fewer than 1,024 variants are padded to fill the context window. This strategy ensures that the model prioritizes the most functionally disruptive variation while maintaining computational efficiency. The resulting input for each gene *g* is a [*N*_max_, 3] tensor, which is processed by the transformer encoder to learn expressive embeddings of the gene’s variant landscape.

### Varformer architecture and optimization

#### Transformer encoding of genetic variation

For a gene *g* and ancestral population *a* ∈ A, let **V**^*g,a*^ = (*v*_1_, …, *v*_*N*_*g,a* ) denote the sequence of observed missense variant tokens, where *N* ^*g,a*^ is the number of tokens associated with gene *g* in population *a*. Each token represents a retained annotation of a DNA variant and is indexed by *i*. Each token *v*_*i*_ is represented by its AlphaMissense pathogenicity score *α*_*i*_ ∈ [0, 1], model-position index 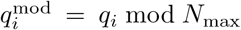 mod *N*_max_ derived from the reported amino acid coordinate *q*_*i*_, and mutation type *m*_*i*_ representing the corresponding amino acid substitution (REF → ALT).

As described above, variants are ranked by decreasing AlphaMissense score and truncated or padded to a fixed sequence length *N*_max_ = 1, 024. We denote the resulting fixed-length sequence by 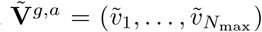 and its associated padding mask by 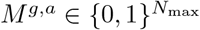, where masked positions correspond to padding.

For each non-padded variant 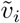, the model computes an initial embedding **e**_*i*_ by concatenating a learned projection of its pathogenicity score with an embedding of its mutation type and adding a positional encoding derived from its sharded model-position index:

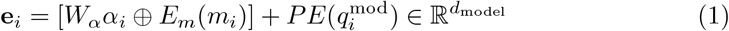

where 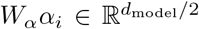 is the learned projection of the scalar pathogenicity score, 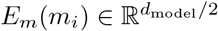 is the learned mutation-type embedding, and 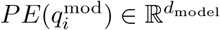 is a sinusoidal positional encoding indexed by the sharded model position 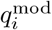 [48].

The resulting variant embeddings are stacked to form

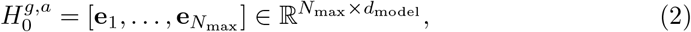

and processed through *L*_*e*_ transformer encoder layers:

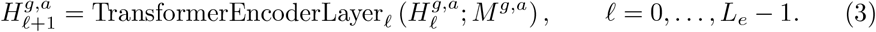

The padding mask *M*^*g,a*^ prevents padded positions from contributing as keys or values during self-attention. This yields a contextualized variant representation 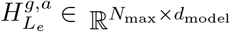, where each variant token is represented in the context of the broader population-specific variant landscape of gene *g*.

#### Projection of gene-level features

In parallel to the PVC branch, the two components of the GC module, Target tractability and safety features 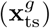 and Target function and expression features 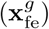, are independently projected into learned gene-level representations:

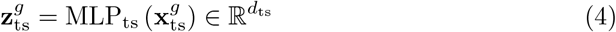

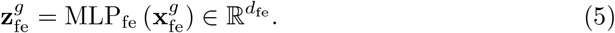

The outputs are concatenated to form a unified gene-level context vector:

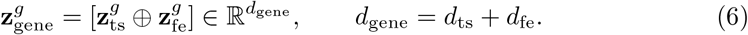

#### Gene and variant prioritization

The contextual variant embeddings get routed to two tracks in parallel. A shared linear head scores individual variant tokens directly to enable variant prioritization within and across genes, while gene-to-variant cross-attention combines the variant embeddings with gene features to produce a gene-level representation that can be used for downstream target prioritization.

Each non-padded row of 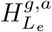 is a contextual variant-token embedding. Writing the embedding of token *i* from model instance *j* as 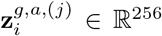, a shared linear head produces a scalar logit and the corresponding vector of token scores:

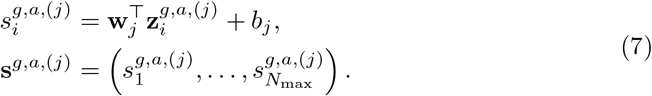

Here *j* indexes the five independently trained instances. The head parameters are shared across all tokens and genes within an instance. The seed index is omitted when describing a single instance, giving **s**^*g,a*^ for its head output. The head predicts scores the from the variant tokens before gene–variant cross-attention, so gene-feature tokens do not directly enter the variant prediction. The gene and variant-level objectives do however update the encoder during joint training. Each token score therefore depends on its AlphaMissense value, substitution identity and model position contextualized by the retained variants in its gene. Higher scores give higher priority for follow-up. Note that the produced logit is not a calibrated probability of pathogenicity. Padding is masked and only retained observed missense tokens are scored.

To fuse the gene-level and population-specific variant representations, Varformer employs a gene-to-variant cross-attention mechanism. The gene context vector 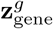 is projected into a query representation, while the transformer-encoded variant sequence 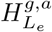 is projected into key and value representations:

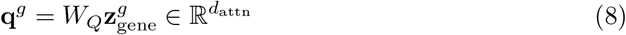

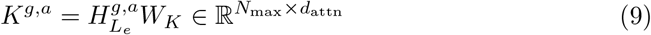

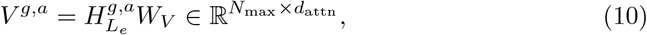

where *W*_*Q*_, *W*_*K*_, and *W*_*V*_ are learned projections into the *d*_attn_-dimensional cross-attention space. The variant-conditioned gene representation is then computed as:

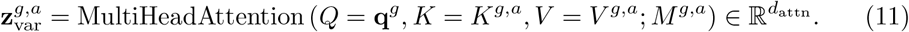

The same padding mask *M* ^*g,a*^ excludes padded variant positions from the cross-attention operation. By conditioning attention on the gene-level context, the contribution of each variant to 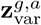 depends on both its contextualized population-specific representation and the gene-level features relevant to therapeutic development.

The final multimodal representation is obtained by concatenating the gene-level and variant-conditioned gene representations:

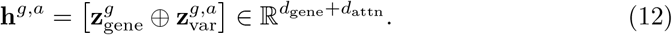

This representation is passed through an *L*_*c*_-layer classification head to produce the final prioritization score:

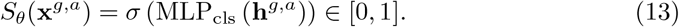

Here, *σ* denotes the sigmoid activation function, *θ* denotes the learned model parameters, and

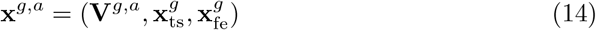

represents the multimodal input features for gene *g* in population *a*, comprising the population-specific variant sequence **V**^*g,a*^ and the gene-level tractability, safety, functional, and expression features. The resulting *S*_*θ*_(**x**^*g,a*^) is a prioritization score rather than a calibrated probability of clinical trial success, as discussed below.

For reporting, *v* identifies a DNA variant by chromosome, position, reference and alternate allele. The score 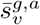 is the arithmetic mean across the five seeds and retained annotations of *v* within gene *g*. Omitting *g*, 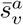averages across all retained annotations of that DNA variant, including those associated with overlapping genes, with equal weight per annotation and seed. Thus each DNA variant receives one reported score. Since we exclusively evaluate variant prioritization for the SAS cohort, we suppress the population superscript and write 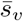.

#### Training labels

We define positive drug targets as genes with clinical-trial-validated therapeutic potential, and we compiled them from three sources. The first source is the Open Targets Platform [17], from which we took genes that have reached at least Phase III clinical trials or approval, corresponding to a `maxClinicalTrialPhase` ≥ 0.75. The second source is Citeline’s Pharmaprojects database, from which we took the targets of drugs with a highest development status of launched, registered, pre-registration or Phase III. The third source is the list of FDA-approved drug targets of the Human Protein Atlas [49]. A gene is assigned a positive label if it appears in at least one source. In total, 1,614 positive targets were present in at least one of the four ancestral population cohorts (1,613 in AFR, 1,612 in AMR, 1,614 in NFE and 1,464 in SAS), of which 1,464 were present in all four.

The remaining genes in the exome are left unlabeled. Within our positive-unlabeled (PU) learning framework, these are treated as having an unknown therapeutic status rather than being definitively non-therapeutic. The model’s objective is to identify candidates within this unlabeled pool that exhibit biological and genetic signatures similar to those of established successful targets.

Variant supervision uses the maximum available adjusted allele frequency across the gnomAD AFR, AMR, NFE and SAS groups. Variants with maximum frequency below 10^−3^ receive label 1, while those with frequency from 10^−3^ to below 0.999 receive label 0. This objective uses low population allele frequency as a proxy for selective constraint, rather than curated pathogenicity labels. Allele frequency is used for supervision and is not an input feature of the variant encoder. Unmatched variants and frequencies outside these ranges are masked from the variant loss. A deterministic 15% gene partition withholds variant labels for evaluation of the frequency proxy. The gene labels and input tokens remain available to the gene objective.

#### Model training

The joint training objective combines gene-level and variant-level supervision:

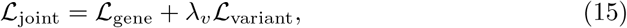

where *λ*_*v*_ controls the contribution of variant supervision and is set to 0.1 for the reported jointly trained models.

The gene loss, ℒ_gene_ ≡ ℒ_nnPU_, is the non-negative positive-unlabeled objective [13]. Let 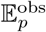 denote expectation over observed positive genes, *p*(**x** | *Y* = +1, *O* = +1), and E_*u*_ expectation over the marginal input distribution *p*(**x**). The class prior *π* is estimated separately for each population as the fraction of positively labeled genes among all genes considered in that cohort. With logarithmic loss, the objective used with observed positives is

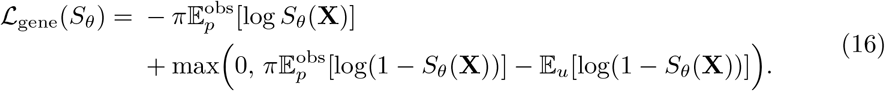

Expectations are estimated by sample averages. The first term penalizes low scores for observed positives; the remaining term estimates negative-class risk from the positive and unlabeled data and clips this estimate at zero. Because observed positives are selected clinical targets, this objective uses their observed distribution rather than the unknown distribution of all true positives. The resulting score is therefore interpreted as a prioritization score, not a calibrated probability of clinical success. The intermediate risk identities, density-ratio interpretation and their assumptions are given in Supplementary Note 6.

For the eligible variant tokens ℳ in a training batch, the variant loss is binary cross-entropy:

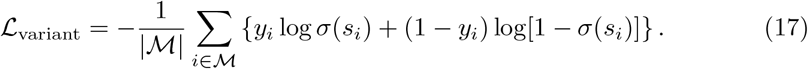

Here *i* indexes eligible tokens in the batch, *s*_*i*_ is an individual model’s raw token logit, *y*_*i*_ is its frequency-derived label and *σ* is the logistic function. Gene, population and seed indices are suppressed in this loss expression. Binary cross-entropy is averaged over eligible tokens without class weighting. Batches without eligible tokens contribute only the gene loss.

All reported models are trained with ℒ_joint_. For each of the four populations, we train five instances with seeds 13, 23, 42, 85 and 482. Training uses AdamW with a batch size of 128, weight decay of 3×10^−4^ and dropout of 0.3. The learning rate follows cosine annealing with warm restarts between 10^−4^ and 10^−5^, with a cycle length of 200 optimizer steps. Each instance is trained for 100 epochs with 32-bit precision, gradient clipping at 1.0 and a maximum norm of 20 for the mutation embeddings. Each instance is selected at the earliest epoch attaining maximum validation gene Spearman correlation, computed once over all validation genes. The hyperparameter search is described in Supplementary Note 8.

For gene classification, the decision threshold is the (1 − *π*) quantile of validation scores, so that the fraction above the threshold follows the estimated class prior. The threshold derivation is provided in Supplementary Note 6.

### Benchmarking setup and evaluation strategy

In addition to Varformer, we benchmarked three alternative models: DrugnomeAI, a logistic regression variant of Varformer, and a random baseline.

DrugnomeAI is an ensemble machine learning framework developed to estimate the druggability of protein-coding genes across the human exome [25]. It integrates gene-level features from heterogeneous data sources and applies a stochastic semi-supervised learning strategy to rank genes by their likelihood of being viable drug targets. The published model is trained on curated sets of druggable genes from the Pharos [50] and Triage [28] repositories. To align DrugnomeAI with our objective, we instead trained it on the positive labels used for Varformer, after removing all benchmark targets (1,556 genes, of which 1,540 are in the DrugnomeAI gene set). We used the released code with the Gradient Boosting classifier, the best-performing configuration reported by Raies et al. [25], and trained one model for each of the five seeds. The default feature set of DrugnomeAI encompasses 323 features. Of these, 60 record drug–gene interactions directly: the number of DGIdb interaction types and 59 chemical–gene interaction features from CTD. Such records exist largely because a gene is already the target of a drug, so they encode the target status that the benchmarks test. The DrugnomeAI model reported in the main text excludes these 60 features. It uses the remaining 263 features, which comprise 165 pathway membership features distributed by CTD, protein family features from InterPro [51], protein–protein interaction network features, gene expression, and other gene-level features. Results for the default feature set and for a feature set that also excludes the pathway features are reported in Supplementary Note 9. DrugnomeAI scores were mapped from gene symbols to Ensembl gene identifiers, and the same DrugnomeAI model of a given seed was evaluated on the benchmark sets of all four populations.

The Varformer logistic regression (Varformer-LR) model serves as a feature-aggregation baseline. Instead of the full transformer architecture, we construct fixed-length gene-level representations from the variable-length variant input. While the gene-level features (GC) are included directly, the PVC features are summarized into 10 descriptive statistics designed to capture the burden, severity, and spatial distribution of mutations for a given gene. Specifically, we compute the number of variants, the mean, standard deviation, maximum, and minimum of pathogenicity scores, as well as the mean and standard deviation of the amino acid positions affected. To quantify heterogeneity in the variant landscape, we compute the *position diversity ratio*, defined as ^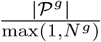^, where *P*_g_ is the set of unique variant positions for gene *g* and *N* ^*g*^ is the total number of variants for gene *g*. The position diversity ratio captures how dispersed the variants are across the protein sequence. Values near 1 suggest broad distribution, whereas lower values indicate concentration at a few positions, possibly reflecting functional hotspots. Additionally, to reflect spatial clustering, we compute the mean and standard deviation of the distances between consecutive variant positions, based on the sorted list of observed amino acid coordinates. This logistic regression baseline provides a point of comparison by summarizing variant information through hand-crafted statistical features. By benchmarking against this baseline, we can assess the added value of the transformer-based architecture and evaluate the extent to which Varformer autonomously learns relevant biological representations from raw sequence data, rather than relying on explicit feature engineering.

Finally, we include a random baseline that classifies each gene as likely to succeed in clinical trials with 50% probability, and as unlikely with 50% probability. This baseline provides a minimal reference point for evaluating predictive performance, representing the expected behavior of a non-informative classifier under uniform class assumptions.

We report three evaluation metrics. Because target prioritization is in practice concerned with the top of the ranked list, we first report precision@20: the proportion of the 20 highest-scoring genes that are true positives. Ranking quality over the complete benchmark set is measured with the Spearman rank correlation *ρ* between predicted scores and ground-truth labels. Classification quality is measured with the F1 score obtained when the *n*_+_ highest-scoring genes are called positive, where *n*_+_ is the number of positives in the benchmark set. For every model, each metric was computed once per run from the predictions for the complete benchmark set, restricted to the genes scored by all three models. All models were evaluated with five random seeds across the four ancestral cohorts, giving 20 runs per reported value. The seed governs both model initialization and the train/validation partition, and identical seeds were used for every model so that all are evaluated on the same partitions. We compared Varformer with DrugnomeAI and baselines over the 20 matched runs with a two-sided sign-flip randomization test (100,000 draws). Confidence intervals were obtained with a hierarchical bootstrap that resamples populations and then seeds within populations (50,000 draws). *P* values were adjusted with the Benjamini–Hochberg procedure across the three benchmarks and the two competing models, separately for each metric.

### Time-stratified prospective validation

The prospective analysis compared the Open Targets target-prioritization feature tables used for training (release 23.06; 29 June 2023) with release 26.06. The baseline cohort comprised genes with `maxClinicalTrialPhase` equal to 0.25 or 0.50 (Phase I or II) in release 23.06 that remained unlabeled during Varformer training. A positive prospective outcome was defined as a transition of the corresponding Open Targets feature to `maxClinicalStage` equal to 0.70 or 1.00 (Phase III or approval) in release 26.06. Eleven *IFNA* genes that map to one sifalimumab program were excluded. Confirmed negative outcomes were identified from trial records and were restricted to directly target-attributable Phase II trials for which the registry record or reported results documented termination for efficacy or safety, or failure of the primary efficacy endpoint. One of the three targets with a confirmed negative outcome, *TREM2*, has no clinical phase annotation in release 23.06 and was included on the basis of its registered Phase II trial. Ongoing, completed-without-results, administratively terminated and otherwise unresolved programs were not assigned negative labels.

Prospective scoring reused the cross-population transformation (Cross-population de-risking via rank-normalized scores). A gene was eligible when it had five-seed ensemble predictions in at least two populations. Its prospective score was the mean of the available population-specific rank-normalized scores 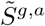. Because the distribution of this mean differs according to the number of populations with available predictions, classification thresholds were estimated separately for genes with predictions in two, three, and four populations. Within each of these three groups, the threshold was the (1 − *π*) quantile of the scores for training-unlabeled genes, with *π* = 0.089, the class prior of the SAS training data (the class priors of the four populations range from 0.082 to 0.089). On the percentile scale, the thresholds were 48.50 for genes with predictions in two populations, 81.18 for genes with predictions in three populations, and 85.30 for genes with predictions in all four populations. The resulting analysis then considers clinical trial events on a target level (Supplementary Table 4).

### Cross-population de-risking via rank-normalized scores

The raw Varformer score *S*_*θ*_(**x**^*g,a*^) represents a proxy for clinical trial success, but it is distinctly not the true probability of clinical trial success *p*(*Y* = +1 | **X** = **x**^*g,a*^) for gene *g* in population *a*. This is because our positive labels (targets reaching Phase III or higher) suffer from a strong selection bias, i.e. targets are not labeled at random but are instead the result of complex filtering based on numerous factors, for example, pre-existing research bias, economic considerations, assessments of druggability and clinical trial quality. This selection bias makes the true posterior *p*(*Y* = +1 | **X** = **x**) non-identifiable [14].

Instead of learning *p*(*Y* = +1 | **X** = **x**) directly, we aim to learn properties of *p*(*Y* = +1 | **X** = **x**) that we can access via a set of observations *O* ∈ {0, +1}, an approach that is termed *partial identification* in statistics and economics [52, 53]. Specifically, Varformer was trained to estimate the density ratio 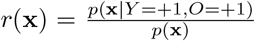 . We assume *invariance of order* : that the observation probability *p*(*O* = +1 | **X** = **x**^*g,a*^) and the true success probability *p*(*Y* = +1 | **X** = **x**^*g,a*^) induce the same ordering over genes. Under this assumption, the learned score *S*_*θ*_(**x**^*g,a*^), which is proportional to *r*(**x**^*g,a*^) within population *a* up to boundary clipping, is order-preserving with the unlearnable *p*(*Y* = +1 | **X** = **x**). Therefore, the meaningful signal from our model is not the absolute value of *S*_*θ*_(**x**^*g,a*^), but rather its relative rank within population *a*.

For the cross-population analysis, scores were first averaged over the five independently trained models within each population. Denoting the parameters of the model trained for population *a* with seed *s* by *θ*_*a,s*_, the ensemble score for gene *g* is

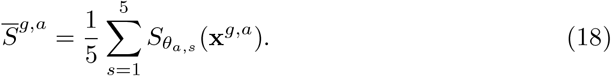

Let *R*_*a*_(*g*) denote the descending rank of 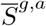 among the *N*_*a*_ genes scored in population *a*, with rank 1 assigned to the highest score and tied scores assigned their average rank. We then apply the following non-parametric transformation:

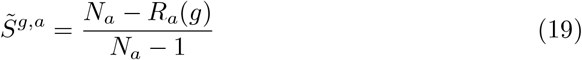

The resulting 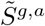 lies in [0, 1], with larger values indicating stronger prioritization relative to other genes scored in the same population.

Only genes scored in all four populations were retained, and targets already carrying a positive training label were excluded. For gene *g*, we summarized the four normalized scores by their mean and sample SD. Figure 4A expresses these values as percentiles where the 1,000 candidates with the highest mean percentile form the overview, and the ten with the highest mean percentile are highlighted. If two genes have the same within-population score, they are assigned the average of their occupied ranks. No ties occurred among these 1,000 candidates. The SD is not used as a second ranking criterion, but measures whether high prioritization is maintained across the four population-specific genetic contexts. We describe candidates combining a high mean percentile with low SD as computationally de-risked against population-level genetic variation.

The Open Targets features are provided on bounded, directionally aligned scales, such that higher values consistently indicate more favorable target properties. Because their empirical distributions differ across features, for Figure 4B we converted each feature to a z-score over the union of genes with available Open Targets features across the four cohort tables. This transformation preserves the directionality of the original scores while expressing each value relative to its feature-specific genomic background. Zero denotes the genome-wide mean, and positive or negative values indicate values above or below that mean, respectively, in units of standard deviation. For Figure 4C, the four most frequent biological process terms and four most frequent molecular function terms among the top ten candidates were selected, and membership was displayed as a binary matrix.

Top-*K* stability was calculated independently within each population. For each seed and each value of *K*, we determined the fraction of that seed’s top-*K* genes that belonged to genes occurring in the top-*K* set of at least two or at least three of the five seeds, and averaged this fraction across seeds. The random reference conditions on a randomly selected gene being present in the seed’s top-*K* set and evaluates the corresponding binomial tail over the other four seeds.

### Evaluation of variant prioritization

#### ClinVar variant classification

We evaluated discrimination between ClinVar pathogenic/likely pathogenic and benign/likely benign variants observed in G&H, using one row per scoreable genomic variant. ClinVar labels were taken from the clinical significance annotation (`CLIN SIG`) of the G&H variant annotation, which was generated with Ensembl VEP (release 111; GRCh38). This annotation does not include the ClinVar review status, so we applied no filter on the level of evidence and included assertions of any review status. A variant was labeled pathogenic if all reported significance terms were pathogenic or likely pathogenic, and benign if all terms were benign or likely benign. Variants with uncertain or conflicting terms, with both pathogenic and benign terms, or without annotation were excluded. AUROC and average precision were calculated separately for rare variants (cohort AF *<* 10^−3^) and across all frequencies, with all methods evaluated on the same model-covered variants within each set. Cohort allele frequency (AF) is the frequency of the alternate allele among G&H participants. We define allele rarity as the negated cohort AF, so that rarer variants receive higher scores. Allele rarity serves as a frequency-only baseline and as a component of the score combinations. Before combining scores, we converted each of the evaluated methods’ scores to percentile ranks within the evaluated variant set, placing all methods on a comparable scale. Variants with identical scores shared the average of the ranks they occupied. Variants with missing model scores were placed at the bottom of that method’s ranking. We averaged these percentiles with equal weights, where, in combination labels, AF denotes allele rarity and AM denotes AlphaMissense. A control assigned two-thirds weight to the AlphaMissense percentile and one-third to the allele-rarity percentile, matching rarity’s weight in the three-way score. No weights were fitted to the evaluation labels. Paired differences were evaluated using 1,000 bootstrap draws resampling genes and retaining their variants together, with the same sampled genes used for both methods and percentile scores held fixed. We report pointwise 95% percentile intervals for these comparisons (Supplementary Note 4 and Supplementary Table 7).

#### Individual-level variant prioritization

Individual-level evaluation used the G&H 55k exome release in its trusted research environment (TRE), with GRCh38 DNA-variant keys and missense annotations on MANE transcripts [54]. The candidate participant pool comprised 3,866 KING degree-3 unrelated individuals absent from the 44k callset used to construct our evaluation set. This excludes those individuals from the input cohort used to train Varformer, but does not imply that their genes or variants were completely unseen. Five-seed mean token scores sharing the same genomic variant identifier were averaged to obtain 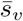.

We describe a variant as carried by a participant if the participant’s genotype contains at least one copy of the alternate allele, in the heterozygous or homozygous state. We refer to such a participant as a carrier; the term does not imply a recessive carrier state or any disease status. For each participant, the candidate variants were all carried missense variants with cohort AF *<* 10^−3^. A candidate variant classified as pathogenic or likely pathogenic in ClinVar was considered a known pathogenic variant. Conflicting classifications were excluded from this definition. Each eligible carrier–variant pair formed a case, so a participant carrying several known pathogenic variants could contribute multiple cases. Cases required at least 20 scoreable variant candidates. Unannotated variants remained candidates and were not labeled benign. The task measures recovery of a known ClinVar annotation within a carrier’s candidate list; it does not establish that the variant caused disease or phenotype in that carrier.

All methods ranked the same 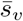-covered candidates. We compared standalone 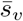, AlphaMissense [55], REVEL [31], CADD [56], allele rarity and a broadcast Varformer gene score that assigns the same score to each variant in a gene. Recall at *k* measures the proportion of evaluated individual–variant pairs for which the known pathogenic variant appears among that individual’s top *k* ranked candidates. We evaluated *k* = 1, 5, 10, 20, 50, and 100. We estimated 95% confidence intervals for differences in recall using 10,000 bootstrap samples. We resampled known pathogenic variants, keeping all carriers of each variant together, and evaluated both methods on the same samples. These intervals are pointwise across the evaluated methods and cutoffs. Additional analyses assessed whether the results changed when restricting candidate lists to variants scored by every method, giving each distinct known pathogenic variant equal weight regardless of its number of carriers, or evaluating known pathogenic variants in genes excluded from variant-level supervision during training. We also tested combinations of scores by averaging their percentile ranks within each individual’s candidate list. These analyses are reported in Supplementary Note 4.

#### Association of variant scores with carrier phenotypes

Our phenotype cohort comprised the 14,698 KING degree-3 unrelated participants in the G&H 55k exome release. Each trait analysis included the participants with a measurement of that trait. Before any trait data were linked to genotypes, we fixed a panel of 15 gene–trait pairs. Each pair consists of a gene with an established effect on a routinely measured quantitative trait and the expected direction of effect of a damaging variant (Supplementary Table 8). No pair was added, removed or changed in direction after the trait data were examined. Qualifying carriers had exactly one rare variant in the gene that was missense in its MANE transcript. Carriers of multiple qualifying variants, unscored variants or annotated loss-of-function variants in that gene were excluded. We regressed median trait measurements on age at measurement, age squared, sex, and ten ancestry principal components, with medication indicators for lipid traits and HbA1c. Residuals were rank-inverse-normal transformed and multiplied by the gene’s hypothesized effect direction. The collapsed DNA-variant scores 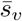 were converted to within-gene percentiles over qualifying variants, so carriers of the same variant shared a score percentile.

We fitted pooled score–trait associations with gene fixed effects in all qualifying carrier observations (subset A) and in observations whose variants lacked a confident ClinVar pathogenic/benign classification (subset B: unannotated, uncertain or conflicting). Here, score–trait association means the fitted regression coefficient relating within-gene variant-score percentile to the adjusted, direction-oriented trait. It measures the estimated trait change per one-unit increase on the 0–1 percentile scale, in rank-inverse-normal trait units. For example, an association of 1 corresponds to an estimated 0.1-unit trait difference for a 0.1 increase in score percentile. Positive values indicate that carriers of higher-ranked variants tend to have greater trait deviations in the expected direction; negative values indicate the opposite, and zero indicates no fitted linear relationship. We used 10,000 bootstrap draws resampling variants within genes and retaining their carriers together. A comparison of pathogenic-variant carriers with non-carriers served as a positive control. We estimated score–trait associations and paired differences from AlphaMissense with bootstrap intervals.

Gene-specific score–trait associations and their mean across genes were exploratory summaries of heterogeneity in the pooled association. For inclusion, a gene required at least 20 carrier observations, ten distinct variants and nonzero score variation in each subset. We fixed these thresholds before running this analysis, as pragmatic minimum requirements for a stable within-gene estimate. Because the score varies between variants and the bootstrap resamples variants, the number of distinct variants limits the precision of each estimate. All 15 genes were retained, including those without sufficient observations for estimation. For each subset, 50,000 variant-bootstrap draws produced pointwise intervals and approximate Bonferroni-adjusted intervals over 60 comparisons (15 genes, two subsets, 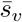 score–trait associations and paired differences from AlphaMissense). Intervals required at least 99% finite bootstrap replicates. For each score and subset separately, we denote the fitted score–trait association for gene *g* by *β*_*g*_ and the set of eligible genes by *G*. The mean score–trait association across genes is 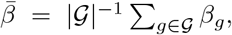, with |*G*| = 10 in each subset. We report pointwise variant-bootstrap intervals for 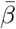. This mean estimates the average association across eligible genes, whereas pooled estimation gives greater influence to genes with more carrier observations and greater score variation. Uncertainty in 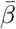 is conditional on the eligible panel and does not account for participants contributing to multiple genes. Leave-one-variant-out analyses assess sensitivity to individual variants.

## Ethics

Genes & Health received a favourable ethical opinion from the NRES Committee London – South East (reference 14/LO/1240) on 16 September 2014. Queen Mary University of London is the sponsor and data controller. All participants gave written informed consent. Individual-level analyses were performed within the Genes & Health trusted research environment, and only aggregate results were exported after review by the Genes & Health data team. The gnomAD, Open Targets and ClinVar data used in this study are publicly available and contain no individual-level information.

## Use of large language models

Between December 2025 and October 2026 we used large language models as coding and writing assistants: Opus 4.6, Opus 4.8, Opus 5 and Opus 5.5 via Claude Code; GPT-5.6 Sol and GPT-6 Astra via Codex. The models helped to draft and edit the manuscript text, to write and run analysis and figure code under the direction of the authors, and to check the consistency of the text with the results. The authors reviewed and edited all model outputs, verified the analyses and the reported values, and take full responsibility for the content of this publication.

## Data availability

Variant data for the NFE, AMR and AFR populations, and the allele frequencies used for variant supervision, are available from gnomAD v4.1 (https://gnomad.broadinstitute.org). Gene-level features and clinical trial status were obtained from Open Targets (release 23.06 for training and release 26.06 for the prospective evaluation; https://platform.opentargets.org). Subcellular location, tissue-level protein expression and UniProt-derived biological process and molecular function annotations were obtained from the Human Protein Atlas version 23 (https://v23.proteinatlas.org), and tissue-level RNA expression from GTEx v10 (https://gtexportal.org). AlphaMissense scores (GRCh38) are available from https://console.cloud.google.com/storage/browser/dm_alphamissense. Positive target labels were derived from Open Targets, Citeline Pharmaprojects and the Human Protein Atlas. The benchmark sets were derived from Pfam (https://www.ebi.ac.uk/interpro/), Pharos v3.18.1 (https://pharos.nih.gov) and Citeline Pharmaprojects (June 2024), which is available under license from Citeline; the genes in each benchmark set are listed in Supplementary Table 3. ClinVar clinical significance (version 2023-06), REVEL (dbNSFP 4.5a) and CADD v1.6 annotations were taken from the Genes & Health variant annotation, generated with Ensembl VEP release 111. Individual-level genetic and phenotype data from Genes & Health are available to approved researchers through application to Genes & Health (https://www.genesandhealth.org) and can be analysed only within its trusted research environment. The Varformer variant scores used in the Genes & Health analyses are available at https://github.com/aaronwtr/Varformer.

## Code availability

Varformer, including the code for model training, inference and variant scoring, is available at https://github.com/aaronwtr/Varformer under the MIT license. The weights of the 25 trained models used in this study are available at https://huggingface.co/aaronw1290/Varformer under the CC BY-NC-SA 4.0 license.

## Acknowledgements

A.W. was supported by the Biotechnology and Biological Sciences Research Council (BBSRC) [BB/X511808/1] through the UKRI AI for Drug Discovery Collaborative Training Partnership, hosted by the Digital Environment Research Institute at Queen Mary University of London, with co-funding from Merck Sharp & Dohme LLC, a subsidiary of Merck & Co., Inc., Rahway, NJ, USA.

Genes & Health has recently been core-funded by Wellcome (WT102627, WT210561), the Medical Research Council (UK) (M009017, MR/X009777/1, MR/X009920/1), Higher Education Funding Council for England Catalyst, Barts Charity (845/1796), Health Data Research UK (for London substantive site), and research delivery support from the NHS National Institute for Health Research Clinical Research Network (North Thames). We acknowledge the support of the National Institute for Health and Care Research Barts Biomedical Research Centre (NIHR203330); a delivery partnership of Barts Health NHS Trust, Queen Mary University of London, St George’s University Hospitals NHS Foundation Trust and St George’s University of London.

Genes & Health has recently been funded by Alnylam Pharmaceuticals, Genomics PLC; and a Life Sciences Industry Consortium of AstraZeneca PLC, Bristol-Myers Squibb Company, GlaxoSmithKline Research and Development Limited, Maze Therapeutics Inc, Merck Sharp & Dohme LLC, Novo Nordisk A/S, Pfizer Inc, Takeda Development Centre Americas Inc.

We thank Social Action for Health, Centre of The Cell, members of our Community Advisory Group, and staff who have recruited and collected data from volunteers. We thank the NIHR National Biosample Centre (UK Biocentre), the Social Genetic & Developmental Psychiatry Centre (King’s College London), Wellcome Sanger Institute, and Broad Institute for sample processing, genotyping, sequencing and variant annotation. This work uses data provided by patients and collected by the NHS as part of their care and support. This research utilised Queen Mary University of London’s Apocrita HPC facility, supported by QMUL Research-IT, http://doi.org/10.5281/zenodo.438045.

We thank: Barts Health NHS Trust, NHS Clinical Commissioning Groups (City and Hackney, Waltham Forest, Tower Hamlets, Newham, Redbridge, Havering, Barking and Dagenham), East London NHS Foundation Trust, Bradford Teaching Hospitals NHS Foundation Trust, Public Health England (especially David Wyllie), Discovery Data Service/Endeavour Health Charitable Trust (especially David Stables), Voror Health Technologies Ltd (especially Sophie Don), NHS England (for what was NHS Digital) – for GDPR-compliant data sharing backed by individual written informed consent.

Most of all we thank all of the volunteers participating in Genes & Health.

## Author contributions

A.W. developed the method, built and trained the model, performed the analyses and wrote the manuscript. C.P.C. contributed to the writing of the manuscript. C.P.C., V.N., W.W. and M.R.B. reviewed the manuscript. A.W., V.N. and M.R.B. conceptualized the project. C.P.C., W.W. and M.R.B. jointly supervised the work.

## Genes & Health Research Team

Eamonn Maher^7^, Shabana Chaudhary^8^, Joseph Gafton^8^, Karen A Hunt^8^, Shapna Hussain^8^, Kamrul Islam^8^, Mohammed Bodrul Mazid^8^, Elizabeth Owor^8^, Jessry Russell^8^, Nishat Safa^8^, John Solly^8^, Marie Spreckley^8^, David A Van Heel^8^, Jan Whalley^8^, Ishevanhu Zengeya^8^, Emily Mantle^8^, Shaheen Akhtar^9^, Samina Ashraf^9^, Dan Mason^9^, John Wright^9^, Daniel MacArthur^10^, Michael Simpson^11^, Richard C Trembath^11^, Gerome Breen^11^, Raymond Chung^11^, Sang Hyuck Lee^11^, Omar Asgar^12^, Joanne Harvey^12^, Karen Tricker^12^, Caroline Winckley^12^, Hanifa Khatun^12^, Amna Asif^12^, Claudia Langenberg^13^, Grainne Colligan^14^, Ceri Durham^14^, Bill Newman^15^, Ahsan Khan^16^, Hilary Martin^17^, Teng Heng^17^, Matt Hurles^17^, Vivek Iyer^17^, Georgios Kalantzis^17^, Vladimir Ovchinnikov^17^, Iaroslav Popov^17^, Klaudia Walter^17^, Panos Deloukas^18^, David Collier^18^, Ana Angel^19^, Saeed Bidi^19^, Fabiola Eto^19^, Sarah Finer^19^, Chris Griffiths^19^, Sam Hodgson^19^, Benjamin M Jacobs^19^, Rohini Mathur^19^, Caroline Morton^19^, Asma Qureshi^19^, Stuart Rison^19^, Annum Salman^19^, Miriam Samuel^19^, Moneeza K Siddiqui^19^, Daniel Stow^19^, Sabina Yasmin^19^, Julia Zöllner^19^, Sheik Dowlut^19^.

^7^Aston University. ^8^Blizard Institute, Queen Mary University of London. ^9^Bradford Teaching Hospitals NHS Foundation Trust. ^10^Garvan Institute. ^11^King’s College London. ^12^Manchester University Hospitals. ^13^Precision Healthcare University Research Institute, Queen Mary University of London. ^14^Social Action for Health (charity). ^15^University of Manchester. ^16^Waltham Forest Council. ^17^Wellcome Sanger Institute. ^18^William Harvey Research Institute, Queen Mary University of London. ^19^Wolfson Institute of Population Health, Queen Mary University of London.

## Supplementary Note 1 Druggability-based cluster analysis of Varformer embeddings

To probe whether Varformer’s gene embeddings encode a notion of druggability that generalizes across populations, we performed an unsupervised cluster analysis on the per-population gene embedding space, supporting the druggability claim in Section 6 and Supplementary Fig. 1.

For each population (SAS, NFE, AFR, AMR), gene-level Varformer embeddings (*d* = 256) were standardized (zero mean, unit variance) and partitioned into two clusters using *k*-means (*k* = 2, `random state` = 42). For all populations, *k*-means was run directly on the 256-dimensional standardized embeddings (`n init` = 20), and UMAP was used only for visualization via a supervised projection onto these cluster labels (`n neighbors` = 500, `min dist` = 1.0, `n epochs` = 1000, `local connectivity` = 2, `repulsion strength` = 0.5).

A gene-level druggability score *s* ∈ {0, 1, 2, 3, 4} was defined as the sum of four Open Targets binary tractability features — `hasLigand`, `hasSmallMoleculeBinder`, `hasPocket`, and `hasHighQualityChemicalProbes` — providing a coarse, interpretable proxy for classical druggability. A gene was deemed *undruggable* if it was unlabeled (i.e. not a known approved target) and had *s* = 0. Druggability distributions in Supplementary Fig. 1 are reported over unlabeled genes only, to avoid conflating the signal with the positive label. Independence of cluster assignment from test-set membership (Pfam / Pharos / Recent) was assessed with a Pearson *χ*^2^ test.

## Supplementary Note 2 Functional characterization of cross-population robust targets

To characterize the leading cross-population candidates, we examined the standardized Open Targets profiles of the five highest-ranked candidates displayed in Figure 4B. All five carry affirmative small-molecule-binder annotations, and four of them (all except *HCN3* ) carry ligand annotations, providing consistent evidence of chemical tractability, although none has a pre-existing annotated druggable pocket. *INSRR* and *TIE1* additionally have high-quality chemical-probe evidence. *INSRR* and *SDHB* carry safety-event annotations, indicating potential safety considerations. *HCN3* has the strongest mouse orthology, whereas *TIE1* and *KCNT1* have the least favorable mouse-knockout scores. *INSRR* and *KCNT1* have favorable tissue-distribution scores, and *SDHB* has the least favorable one.

Because absent annotations do not constitute positive evidence, the profiles indicate where the available evidence supports or cautions against further target development, but do not provide a single measure of druggability. Overall, the candidates share evidence of chemical tractability but differ in the safety, selectivity and experimental-model considerations that would shape their further development, indicating that no single tractability feature explains their consistently high cross-population rankings.

The top ten candidates—*HCN3, INSRR, TIE1, KCNT1, SDHB, CD81, P2RX1, ITPR3, TRPM5*, and *P2RX5* —converge on ion-channel and receptor biology (Fig. 4C). Transport and receptor activity each annotate seven of the ten candidates, ion transport and ion-channel activity six each, and ligand-gated ion-channel activity five; potassium transport, potassium-channel activity, voltage-gated channel activity and tyrosine-protein kinase activity each annotate two. This coherence is driven by purinergic receptors (*P2RX1* and *P2RX5* ), potassium and other cation channels (*HCN3, KCNT1* and *TRPM5* ), intracellular calcium signaling (*ITPR3* ) and two receptor tyrosine kinases (*INSRR* and *TIE1* ).

## Supplementary Note 3 Systematic prospective validation across Phase I/II unlabeled genes

We queried the 332 genes classified at Phase I/II in Open Targets release 23.06 against release 26.06. Of these, 243 belonged to the training-unlabeled pool and had five-seed ensemble predictions in at least two populations. 39 baseline Phase I/II genes moved to Phase III or approval in the updated Open Targets target-prioritization feature. 24 were eligible under our inclusion criteria (training-unlabeled, with predictions for at least two populations). Eleven of these are *IFNA* genes that map to one sifalimumab program and were excluded, which leaves 13 progression targets. We separately reviewed post-cutoff terminal trial records and identified three target-attributable outcomes with explicit efficacy or safety evidence: *DKK1, OSMR* and *TREM2. TREM2* has no clinical phase annotation in release 23.06 and was included on the basis of its registered Phase II trial. Genes with ongoing programs, no reported outcome, or only administrative or recruitment-related termination were left unlabeled and excluded from the validation.

Across the 16 eligible targets, the predictions comprised 10 true positives, one true negative, three false negatives and two false positives, corresponding to an accuracy of 68.8%, sensitivity of 76.9%, specificity of 33.3%, balanced accuracy of 55.1% and an F1 score of 80.0%. With only three confirmed failures, the specificity estimate is imprecise. Using each gene’s distance from the threshold defined for genes with predictions in the same number of populations as a continuous decision value yielded an AUROC of 0.769. Progression events spanned a broad range of cross-population percentiles (median 89.08; range 32.86–98.68), whereas the three confirmed failures ranged from 63.51 to 87.94 (median 87.69). Two of the failed targets, *OSMR* and *TREM2*, were above the four-population threshold. This apparent difference in spread reflects both the small number of confirmed failures and the construction of the cohort. Every evaluated target had already reached Phase I or II by the training cutoff and had therefore passed substantial biological, pharmacological and commercial selection before entering the analysis. Even targets that subsequently failed could consequently retain moderately high Varformer scores. Conversely, some lower-ranked targets progressed because clinical advancement also depends on indication-specific evidence, therapeutic modality, molecule properties and trial design, none of which is fully represented by Varformer’s disease-agnostic target-level features. The wide range among progression events therefore reflects the heterogeneity of routes by which a program can advance, rather than implying that all progressing targets should receive similarly high scores.

Moreover, the distributions of this validation cohort and the classification threshold should be interpreted in light of how we defined our training objective. It was not specifically optimized to separate subsequent progression from failure within this Phase I/II cohort. Instead, the (1 − *π*) quantile rule, with *π*=0.089, designates only the highest-scoring 8.9% of training-unlabeled genes as predicted positives. Applying the same prior-derived rule separately to genes with predictions in two, three or four populations preserves this deliberately selective prioritization criterion in the prospective analysis. A score above the threshold therefore indicates membership in a highly ranked subset relative to genes with the same number of available population predictions, whereas a score below it means only that the gene did not enter this prior-derived subset, not necessarily that clinical failure was predicted. Consistent with this distinction, three targets progressed despite falling below their respective thresholds. Thus, the results support using the threshold to nominate targets with particularly strong model evidence, but not to exclude lower-ranked targets from consideration.

The validation also shows why target-level prioritization can not be equated with the outcome of a particular program as there are too many trial outcome determinants that are not included. Varformer only includes biological-, population-genetic-, and tractability determinants. High-scoring *TLR8, ITGB6* and *WEE1* each reached Phase III while also encountering an efficacy or safety setback in another trial or program (Supplementary Table 4). These mixed histories lay bare the extent to which indication, molecule, regimen and trial design contribute information not resolved by a population-informed target-level score.

### Case Study 1: Multi-modal integration elevates a sparse-variant target (*KDM1A*) subsequently advanced to Phase III

*KDM1A* encodes lysine demethylase 1A (LSD1), a flavin adenine dinucleotide (FAD)-dependent amine oxidase that removes mono- and di-methyl marks from histone H3 lysine 4 (H3K4me1/me2), modulating transcriptional programs essential for hematopoietic progenitor self-renewal and myeloid differentiation [57, 58]. *KDM1A* has attracted sustained interest as a hematological oncology target, with the mechanism-based inhibitor iadademstat (ORY-1001) entering Phase I in acute myeloid leukemia [59] and bomedemstat (IMG-7289/MK-3543) entering Phase I/II in myeloproliferative neoplasms (MPNs) [60]. At the training cutoff, *KDM1A* was assigned a Phase II label in Open Targets and was therefore treated as an unlabeled gene during the training of Varformer.

Only eight *KDM1A* missense variants overlapped between NFE and SAS, of which four met the allele-count and allele-number criteria for enrichment testing. Nevertheless, its mean cross-population percentile was 94.00 (SD 1.68), with population-specific percentiles of 92.00–95.49, placing it above the four-population threshold of 85.30. This high and cross-population robust ranking despite sparse coding-variant representation illustrates the value of Varformer’s multi-modal integration: *KDM1A* carries all major druggability flags (hasPocket = 1, hasSmallMoleculeBinder = 1, hasHighQualityChemicalProbes = 1), features strongly enriched among training-positive genes. Gene ontology annotations further reinforce this signal, with *KDM1A* annotated to histone demethylation (GO:0032452), chromatin remodeling (GO:0006325), and hematopoietic stem cell differentiation, ontological categories highly enriched among training-positive genes in the hematology space [25].

On 31 December 2023, six months after the training cutoff, the randomized Phase III Shorespan-006 trial began evaluating bomedemstat (MK-3543) against the best available therapy in essential thrombocythemia (NCT06079879) [61]. Bomedemstat acts as a mechanism-based irreversible inhibitor that covalently modifies the FAD cofactor in the *KDM1A* active site, permanently halting demethylase activity [60]. This direct Phase III advancement provides prospective support for Varformer’s prioritization of *KDM1A* and illustrates how multi-modal integration can elevate a clinically actionable target that approaches relying solely on population-genetic variant burden would undervalue.

### Case Study 2: A human-genetics-supported regulator of iron homeostasis (*TMPRSS6* ) subsequently advanced to Phase III

*TMPRSS6* encodes matriptase-2, a liver-expressed transmembrane serine protease that suppresses the iron-regulatory hormone hepcidin and thereby promotes systemic iron availability. Human genetics provides direct functional evidence for this pathway: biallelic loss-of-function variants cause iron-refractory iron-deficiency anemia, while the common nonsynonymous variant rs855791 is associated with serum iron, transferrin saturation and erythrocyte volume [62, 63]. At the training cutoff, sapablursen, a liver-directed antisense oligonucleotide that reduces *TMPRSS6* expression, was in Phase II and *TMPRSS6* therefore remained in the training-unlabeled set.

Shared SAS–NFE variants contained 99 *TMPRSS6* missense variants, of which 74 met the criteria for enrichment testing. Its mean cross-population percentile was 88.24 (SD 1.51), with population-specific percentiles of 85.98–89.09, placing the gene above the four-population threshold of 85.30. The agreement across population-specific variation profiles is complemented by the external human-genetic evidence, as both rare severe alleles and a common coding allele alter the same iron-regulatory biology that is therapeutically engaged by suppressing *TMPRSS6*.

On 25 May 2026, sapablursen entered the recruiting Phase III INTREPID trial in polycythemia vera (NCT07429266) [64]. Sapablursen directly suppresses *TMPRSS6*, increasing hepcidin and restricting the iron available for erythropoiesis. This case shows a prioritized gene product is itself engaged by the advancing therapeutic modality. It also illustrates a scenario in which all of Varformer’s contributing modalities are crucial: population-specific missense information is evaluated together with gene-level functional and tractability features, backed up by evidence from the human genetic record.

### Case Study 3: A high-profile target (*TREM2* ) that failed Phase II despite a score above the threshold

*TREM2* encodes triggering receptor expressed on myeloid cells 2, a single-pass type I transmembrane receptor expressed predominantly on microglia in the central nervous system [65]. In 2013, rare coding variants in *TREM2* (p.R47H and p.R62H) were independently identified as moderate-penetrance risk alleles for late-onset Alzheimer’s disease (AD), generating sustained interest in *TREM2* as a microglial activation target for neurodegeneration [66, 67]. At the training cutoff, AL002, an anti-TREM2 agonist monoclonal antibody co-developed by Alector and AbbVie, was in Phase II in the INVOKE-2 trial (NCT04592874) in early symptomatic AD. *TREM2* carries no clinical phase annotation in Open Targets release 23.06 and was unlabeled during Varformer training.

The intersection of SAS–NFE variants contained 30 *TREM2* missense variants, of which 26 met the enrichment-testing criteria. Its mean cross-population percentile was 87.69 (SD 0.57), with population-specific percentiles of 86.86–88.07. This placed *TREM2* above the four-population threshold of 85.30 by a small margin. Its Open Targets profile nevertheless offered limited evidence of classical chemical tractability: *TREM2* lacks small-molecule-binder and druggable-pocket annotations, and its biology is concentrated in microglial innate-immune signaling.

On 25 November 2024, Alector announced that the Phase II INVOKE-2 trial had failed to demonstrate clinical benefit in early AD: no treatment effect was observed on cognitive decline, amyloid PET, or established AD fluid biomarkers, despite pharmacodynamic evidence of target engagement measured by elevated soluble TREM2 in cerebrospinal fluid [68]. The above-threshold *TREM2* prediction is therefore a false positive in our eligible validation cohort. The case illustrates the limit of a gene-level score: it reflects the plausibility of a gene as a target, which the human genetic evidence for *TREM2* supports, and does not anticipate the outcome of a specific molecule, indication and trial.

## Supplementary Note 4 Variant prioritization benchmark coverage and classification

The G&H retrieval score export contains 2,264,706 token rows across 15,725 genes, corresponding to 1,299,645 distinct GRCh38 DNA-variant keys. Five-seed mean token scores were averaged across retained annotations of each DNA variant to obtain 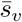before joining genotypes; this includes annotations associated with overlapping genes. The population superscript is omitted for these SAS scores. Predictor annotations with multiple values used their maximum, whereas model scores used their mean; this difference is part of the evaluated implementation. The 55k-release annotation did not provide ClinVar review tiers; known pathogenic variants were therefore selected by clinical significance without an expert-panel restriction. A known pathogenic variant required all nonempty clinical-significance terms to agree on pathogenic or likely pathogenic status. Uncertain and conflicting annotations were excluded from the known pathogenic variant set, not from the candidate background.

Model scores covered 66.79% of carried rare variants and 72.93% of known pathogenic variants. Coverage was assessed before dropping unscored candidates. All reported retrieval curves consequently describe the model-covered subset and cannot establish whole-exome retrieval sensitivity. The complete-case comparator sensitivity retained 1,060 carrier–variant pairs and 547 distinct known pathogenic variants; 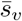 and AlphaMissense top-20 recall were 59.7% and 55.9%, respectively, while REVEL reached 78.6%. Among known pathogenic variants in genes excluded from variant-level supervision (153 carrier–variant pairs, 79 distinct variants), 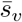 did not show a clear advantage: top-10 recall was 32.7% versus 36.0% for AlphaMissense, with a paired difference interval of −11.5 to 4.4 percentage points. Bootstrap resampling kept all carriers of each known pathogenic variant together; a sensitivity analysis gave each distinct known pathogenic variant equal weight regardless of its carrier count. These comparisons are exploratory. Their full tables appear in Supplementary Data 1; the standalone and combination retrieval results are in Supplementary Table 5.

Known-label discrimination is a separate task: distinguish ClinVar pathogenic/-likely pathogenic from benign/likely benign DNA variants, excluding unannotated, uncertain and conflicting labels. Within the G&H rare-variant subset, 2,477 pathogenic and 15,580 benign scoreable variants were analyzed; 10,833 labeled variants lacked a model score. Across all frequencies, the corresponding numbers were 2,521 pathogenic, 25,777 benign and 14,972 unscored labeled variants. We report AUROC and average precision (AP, the reported precision–recall summary) together because their ranking sensitivities and dependence on class balance differ. Gene-bootstrap intervals used 1,000 draws. Standalone 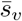 was below AlphaMissense on both metrics in both subsets (Supplementary Table 6).

For secondary combinations, constituent scores were converted to empirical rank percentiles with average ties and missing scores lowest, then averaged with equal weights; no combination weight was fitted on the TRE outcomes. In G&H, AlphaMissense+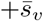 exceeded both individual scores on AUROC and AP in the rare and all-frequency sets. The three-way combination also exceeded a control assigning two-thirds weight to AlphaMissense and one-third to allele rarity, which holds the frequency contribution constant. Removing AF from the three-way score improved both metrics among rare variants but reduced both across all frequencies. Supplementary Table 6 reports all point estimates, including the control; Supplementary Table 7 reports all seven paired comparisons in each set. Intervals used 1,000 paired gene-bootstrap draws, retaining fixed percentile scores, and are pointwise 95% intervals without multiplicity adjustment. The two variant sets overlap and are not independent replications.

In the separate public ClinVar probe-gene evaluation, one row per DNA variant gave 3,314 variants per seed. Here 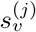 denotes the annotation-averaged DNA-variant score from seed *j*, without averaging predictions across seeds. AUROC and AP were calculated separately for each seed and then averaged. These mean AUROC/AP values were 0.8639/0.5395 for 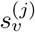, 0.8785/0.5534 for AlphaMissense, 0.9096/0.5999 for 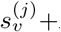, 0.9217/0.6229 for AlphaMissense+AF and 0.9254/0.6784 for the three-way combination. Here AF denotes negated allele frequency, so rarer variants receive higher ranks. The three-way AP difference from AlphaMissense+AF was 0.0564 (95% interval, 0.0283–0.0843), whereas its AUROC difference interval included zero. These are secondary combination results, not gains of standalone 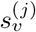. The analysis used mean aggregation across annotations and 400 gene-bootstrap draws per seed; max aggregation was a sensitivity. Repeated annotations are not treated as independent variants.

ClinVar labels are shared evidence across public discrimination, G&H discrimination and retrieval; these analyses are not independent replications of pathogenicity validation. Frequency contributes to clinical classification and to the model’s training proxy. AlphaMissense is itself an input to the variant encoder, and comparator training-data overlap was not resolved in these evaluations. Unannotated candidates ranked above a known pathogenic variant cannot be assumed benign, but neither does high rank establish their pathogenicity.

## Supplementary Note 5 Carrier phenotype results and exploratory gene-specific associations

The fixed gene–trait panel and directions are listed in Supplementary Table 8. Subset A includes all qualifying carriers; subset B excludes confident pathogenic/likely pathogenic and benign/likely benign annotations and retains unannotated, uncertain and conflicting variants. Counts are carrier observations across gene–trait pairs, not necessarily distinct people. The positive control compared pathogenic-variant carriers with non-carriers and gave an oriented trait difference of 1.051 (95% interval, 0.301– 1.621). The pooled 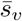 score–trait association in subset B was 0.143 (−0.074 to 0.356), with an interval including zero. Its paired difference in score–trait association from AlphaMissense was −0.106 (−0.261 to 0.042), reported descriptively. All score–trait associations, including AF and score combinations, are retained in Supplementary Table 9.

The exploratory per-gene analysis was motivated by the pooled result and observed heterogeneity. Ten of the 15 genes met the fixed follow-up requirements of at least 20 carrier observations and ten distinct variants in each subset. *GCK, APOA1, LPL, APOA5* and *TSHR* did not meet these requirements. Their comparisons remain in the family of 60 used for interval adjustment. *LDLR* was the only gene with a positive 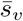 interval excluding zero after this adjustment in both subsets. In subset B, its paired difference from AlphaMissense was −0.060, with a pointwise interval of −0.632 to 0.462. Thus, the positive association does not imply a comparative advantage (Supplementary Fig. 4). The adjusted intervals are approximate bootstrap intervals and do not remove the exploratory nature of the follow-up.

The mean score–trait associations across the ten eligible genes were 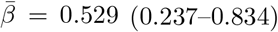 for 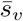 and 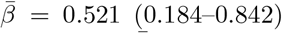 for AlphaMissense in subset A, and 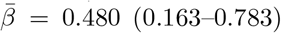 versus 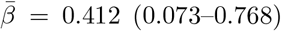 in subset B. Here 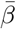 is the arithmetic mean of the fitted gene-specific score–trait associations, calculated separately for each score and subset as defined in Methods (Association of variant scores with carrier phenotypes). The paired differences included zero: 0.008 (−0.275 to 0.354) and 0.068 (−0.248 to 0.407), respectively. These descriptive intervals condition on the eligible panel and do not model dependence from people contributing to multiple genes. They address a different estimand from the carrier-weighted pooled score–trait association.

In subset B, *HBB* and *HNF1A* had positive point differences from AlphaMissense, and leave-one-variant-out difference ranges remained positive (0.335–1.944 and 0.087– 0.264, respectively). Their bootstrap difference intervals nevertheless included zero, so they remain uncertain observations rather than established gains. *APOB* contributed approximately 42% of subset-A carrier observations and had negative score–trait associations for both scores, a gene in which different variants can have opposite trait effects. Its direction was not changed after inspecting the results. Leaving *APOB* out did not establish an advantage over AlphaMissense.

Sensitivity analyses used the 3,866-person input-callset-excluded cohort, excluded medication users rather than adjusting for medication, or used maximum rather than median trait measurements. Results, leave-one-gene-out estimates, incremental models and complete gene-specific and leave-one-variant-out summaries are provided in Supplementary Data 1. Original aggregate outputs and the screenshot-transcribed gene summaries are distinguished in the source provenance. The latter reproduced the original pooled A/B score–trait associations as a continuity check; counts of one to five remain suppressed. No individual-level or per-variant phenotype records are included.

## Supplementary Note 6 Theoretical justification for pseudo classification risk minimization

This note derives the nnPU gene objective, examines the effect of observing only a selected subset of positive genes, and describes the classification threshold used in the training of Varformer.

### From classification risk to nnPU risk

Given an input space *X* and a binary output space *Y* = {+1, −1}, we assume that samples (**X**, *Y* ) are drawn from an unknown joint distribution *p*(**x**, *y*). The class-conditional distributions are defined as:

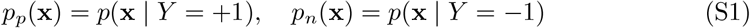

and the marginal distribution of **X** is:

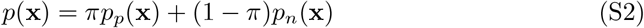

where *π* = *p*(*Y* = +1) is the class prior, estimated separately for each population as the fraction of positively labeled genes among all genes considered in that cohort.

Following the standard PUL setting, the true classification risk for the parameterized decision function *S*_*θ*_ : *X*→ [0, 1] under a loss function *l* : [0, 1] × *Y* → ℝ is given below:

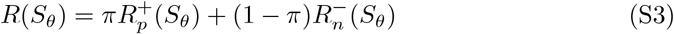

where:

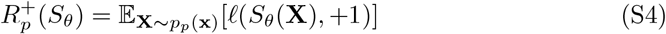

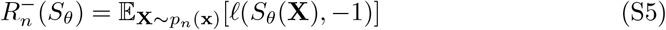

Since negative samples are unavailable, we use the identity *p*_*n*_(**x**) = (*p*(**x**) − *πp*_*p*_(**x**))*/*(1 − *π*) to obtain the unbiased positive-unlabeled risk estimator, as derived in Du Plessis et al. [69]:

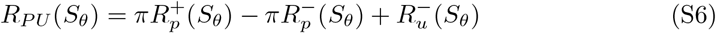

where:

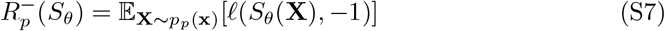

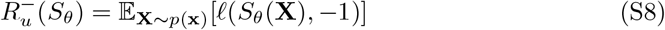

While *R*_*PU*_ (*S*_*θ*_) is an unbiased risk estimator, it can yield negative empirical risk values, leading to overfitting in flexible models. To prevent this, we adopt the non-negative positive-unlabeled risk estimator [13]:

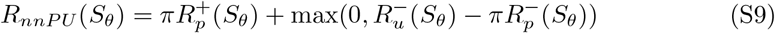

Using logarithmic loss yields the gene objective; Equation (16) in the main text gives its form when observed positives supply the positive-class expectations.

### Observed positives and density-ratio interpretation

We examine the corresponding pseudo classification risk when the positive-class distribution is replaced by *p*(**x** | *Y* = +1, *O* = +1). Under the conditions stated below, its pointwise minimizer is proportional to the observed-positive density ratio *r*(**x**) = *p*(**x** | *Y* = +1, *O* = +1)*/p*(**x**), up to boundary clipping.

Let *S* : *X*→ [*ε*, 1 − *ε*] be a candidate scoring function for the parameterized model *S*_*θ*_, with range clipped to avoid numerical instability, where *ε >* 0 is a small constant that we introduce in order to make the optimization problem well-defined. Define:

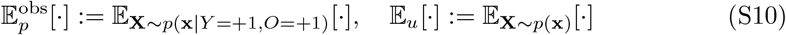

Then the pseudo classification risk becomes:

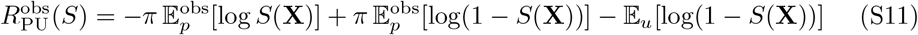

Let 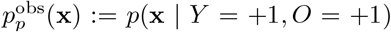 and let *p*(**x**) denote the marginal over **X**. To find our model, we aim to solve the following functional minimization problem.

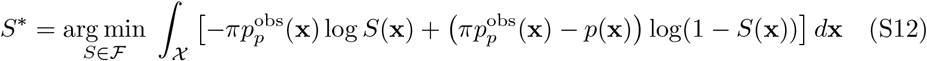

We now analyze the pointwise minimizer of the functional in (S12). We can reduce this minimization problem to a pointwise convex optimization by introducing a scalar variable for each input **x** ∈ *X*. Let

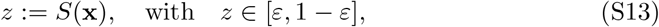

where *S*(**x**) is the model’s predicted probability for the positive class at input **x**, and the interval constraint ensures numerical stability.

We now formulate the resulting scalar optimization problem for each **x**. We can write the risk function for each **x** ∈ *X*as:

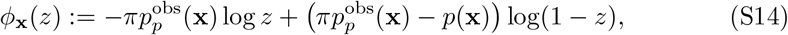

This gives the constrained scalar minimization problem:

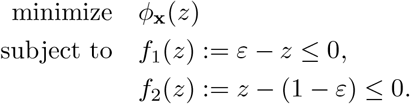

This is a convex optimization problem since *ϕ*_x_(*z*) is convex over the interval *z* ∈ (*ε*, 1 − *ε*), and the box constraints define a convex feasible set. Moreover, the inequality constraints are convex as they are both affine functions of *z*. Here we prove that indeed *ϕ*_x_(*z*) is convex.

#### Theorem 1.

*We define*

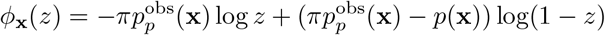

*Suppose that the density ratio satisfies*

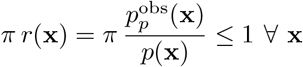

*Then ϕ*_x_(*z*) *is convex on* (*ε*, 1 − *ε*), *where ε >* 0 *is a small positive constant. Proof*. To prove convexity, we need to show that

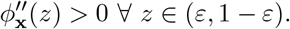

To this end, we compute the second derivative of *ϕ*_x_(*z*) with respect to *z*.

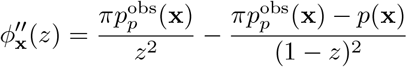

Substituting in the previously defined density ratio *r*(**x**), we get

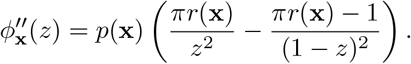

Let us define

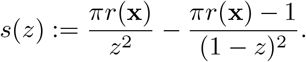

Since *p*(**x**) *>* 0, we have 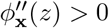 if and only if *s*(*z*) *>* 0.

For *z* ∈ (*ε*, 1 − *ε*) with *ε >* 0 small, both *z*^2^ and (1 − *z*)^2^ are strictly positive and bounded away from zero. Since *π >* 0 and *r*(**x**) *>* 0, we consider two cases:

#### Case 1

If *πr*(**x**) = 1, then

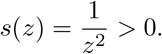

#### Case 2

If *πr*(**x**) *<* 1, then both terms in *s*(*z*) are finite, the numerator of the second term is negative for all *z* ∈ (*ε*, 1 − *ε*), and so we have

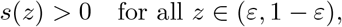

Therefore, 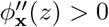 for all *z* ∈ (*ε*, 1 − *ε*), proving that *ϕ*_x_(*z*) is strictly convex on this interval.□

Now, we can apply the Karush-Kuhn-Tucker (KKT) conditions to solve this problem. Define scalar dual variables *λ*_1_ ≥ 0 and *λ*_2_ ≥ 0 for the inequality constraints *f*_1_(*z*) and *f*_2_(*z*) respectively. The Lagrangian is then given by:

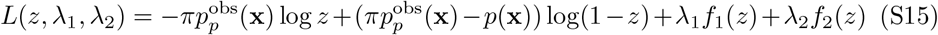

The KKT conditions are:

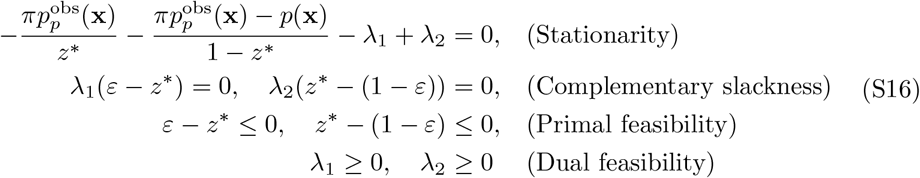

We solve the resulting system case-by-case in order to find the optimal solution on the clipped domain *z* ∈ [*ε*, 1 − *ε*]. We denote this clipped solution as the *optimal decision function*, for which *S*^*^(**x**) = *z*^*^ will hold by construction.

For notational convenience, let us define the following regions:

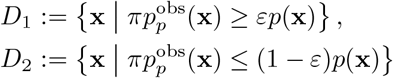

We now analyze each possible KKT solution:

#### Case 1: Interior solution

Suppose *ε < z*^*^ *<* 1 − *ε*. Then both constraints are inactive, so *λ*_1_ = *λ*_2_ = 0. The stationarity condition becomes:

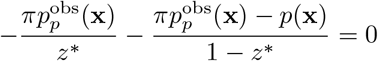

Solving for *z*^*^, we find:

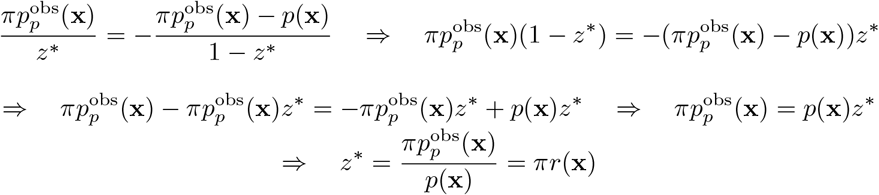

#### Case 2: Left boundary solution

At the *z*^*^ = *ε, λ*_1_ is active while *λ*_2_ is inactive. Plugging this into the stationarity condition then yields:

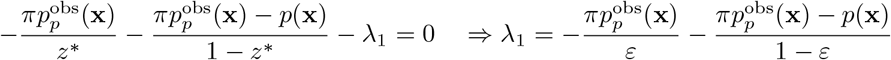

By dual feasibility we must have:

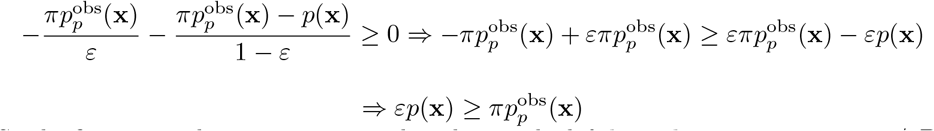

So the first inequality constraint implies that at the left boundary *z*^*^ = *ε* ⇐⇒ **x** ∈*/ D*_1_

#### Case 3: Right boundary solution

At the right boundary we have *z*^*^ = 1 − *ε* which implies that *λ*_2_ is active while *λ*_1_ is inactive. The stationarity condition then yields

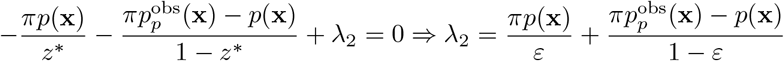

By dual feasibility we must have:

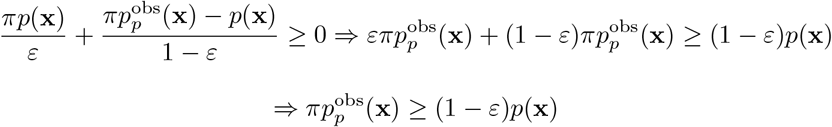

The second inequality constraint thus implies that at the right boundary we must have *z*^*^ = 1 − *ε* ⇐⇒ **x** ∈*/ D*_2_

Combining all three of the above cases, we conclude that the minimizer is given by:

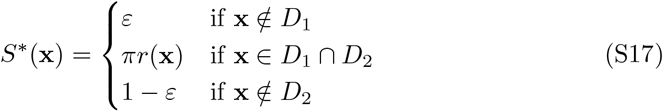

Therefore, minimizing the pseudo classification risk recovers a clipped version of the true density ratio that is almost everywhere equal to *πr*(**x**). Dividing by *π* yields a clipped estimator of *r*(**x**), as desired.

### Classification threshold

Following Kato et al. [14], after estimating *r*(**x**), we determine a classification threshold *τ*_*π*_ by ensuring

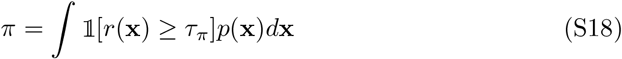

The intuition behind this constraint on *τ*_*π*_ is that we do not want the proportion of positives in our predictions to deviate too much from the class prior *π*. Since this integral cannot be directly computed, we estimate *τ*_*π*_ from a finite validation dataset 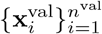 by solving:

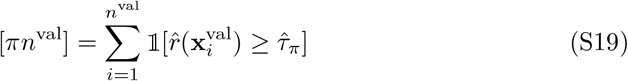

where [*πn*^val^] denotes the nearest integer to *πn*^val^. In practice, this is efficiently approximated using the quantile function:

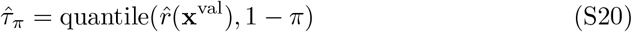

which selects the (1 − *π*)-th quantile of the estimated density ratios. This ensures that the fraction of predicted positive samples aligns with the prior *π*, maintaining class balance.

## Supplementary Note 7 Preprocessing of gene characterization features

To ensure consistency across heterogeneous data sources, the features utilized in the Gene Characterization (GC) module were derived from the Open Targets Target Prioritization dataset [17], which provides a standardized scoring framework for therapeutic targets. Unlike raw genomic metrics, these features are provided as inherently normalized scores within the range [−1, 1].

In this system, a score of +1 indicates maximum evidence supporting a target’s suitability for drug discovery (e.g., the presence of high-quality chemical probes or known druggable binding pockets), while a score of −1 indicates evidence suggesting reduced therapeutic potential or increased risk (e.g., known safety events). Features capturing purely positive evidence, such as ligand availability, are restricted to the [0, 1] interval. This standardized scale allows Varformer to integrate diverse biological signals, ranging from molecular tractability to physiological specificity, without requiring additional normalization or feature-specific weighting.

## Supplementary Note 8 Hyperparameter optimization

Hyperparameter optimization for Varformer was conducted using Optuna [70], employing the Tree-structured Parzen Estimator (TPE) sampler to navigate the parameter space. We performed a search budget of 100 trials, with the objective of maximizing the validation F1 score. To improve search efficiency, trials were pruned using the `PyTorchLightningPruningCallback` if their intermediate performance fell below the median of previous successful trials at the same step.

The search space included both architectural and training hyperparameters. Learning rate followed a cosine annealing schedule [71] with warm restarts, where the initial rate was sampled from {10^−5^, 3 × 10^−5^, 10^−4^} and the final rate was set to either 10% or 1% of the starting value. Weight decay was explored across {3 × 10^−4^, 5 × 10^−4^, 3×10^−3^, 5×10^−3^}. For the Varformer architecture, we evaluated encoder depths (*L*_*e*_) of {2, 3, 4, 6, 8} layers, attention heads (*n*_head_) of {4, 8, 16}, and model dimensions (*d*_model_) of {128, 256, 512, 768, 1024}. The classification head depth (*L*_*c*_) was varied between {2, 4, 6} layers.

The selected configuration consisted of a 3-layer transformer encoder (*L*_*e*_ = 3) with 8 attention heads (*n*_head_ = 8) and an embedding dimension of 256 (*d*_model_ = 256), followed by a 4-layer classification head (*L*_*c*_ = 4). The reported models use the AdamW optimizer with a batch size of 128, a dropout rate of 0.3 and a weight decay of 3×10^−4^. The learning rate follows a cosine annealing schedule with warm restarts between 10^−4^ and 10^−5^, with a cycle length (*T*_0_) of 200 optimizer steps. Each model is trained for 100 epochs with 32-bit precision and gradient clipping at 1.0, and is selected at the earliest epoch attaining maximum validation gene Spearman correlation. Furthermore, rather than using a static classification threshold or a weighted accuracy metric, the optimal decision threshold *τ* was determined dynamically during training as the 1 − *π* quantile of the validation probability distribution, where *π* represents the estimated class prior. This approach ensures that the model’s classification output reflects the expected prevalence of therapeutic targets in the unlabeled exome.

## Supplementary Note 9 Adaptation of DrugnomeAI to prevent label leakage

We use a narrow definition of label leakage: a feature leaks if it directly records that a gene is, or has been, the target of a drug. A measurable property of a gene that correlates with success as a target, such as tractability or tissue expression, is not leakage under this definition.

The default feature set of DrugnomeAI (323 features) contains two groups of features that meet this definition. The first is the number of interaction types that the Drug–Gene Interaction Database (DGIdb) records for a gene. The second consists of 59 features from the Comparative Toxicogenomics Database (CTD) that count chemical– gene interactions, in total or of a given type, for example the chemicals that decrease the activity of the gene product. A gene acquires such records mainly when compounds are developed against it. The remaining 165 CTD features describe membership of KEGG and Reactome pathways and do not record interactions with drugs.

The benchmark sets show the effect directly. At least one DGIdb interaction type is recorded for all Pfam positives and 93% of the Recently Approved positives, compared with 8% of the benchmark negatives. Used alone as a score, the DGIdb feature separates benchmark positives from negatives with an AUROC above 0.99 on Pfam, 0.96 on Recently Approved and 0.67 on Pharos, and the CTD count of unique interactions reaches 0.90, 0.82 and 0.50, respectively (means over the 20 benchmark sets). The Pharos targets have potent ligands but no approved drug, which is consistent with the weaker separation on that benchmark.

We therefore trained DrugnomeAI without these 60 features, which is the configuration reported in the main text (263 features). For comparison, we trained two further configurations on identical labels and seeds: with the default features, and without DGIdb and all 224 CTD features, which also removes the 165 pathway features. With the default features, DrugnomeAI exceeded Varformer on all three metrics on Pfam and Recently Approved (adjusted *q <* 0.05); on Pharos, Varformer had the higher Spearman correlation and the other two comparisons were inconclusive. Removing the 60 interaction features lowered the precision@20 of DrugnomeAI from 0.97 to 0.91 on Pfam, from 0.95 to 0.84 on Recently Approved and from 0.53 to 0.39 on Pharos. In this configuration, Varformer exceeded DrugnomeAI in four of the nine comparisons (precision@20 on Pfam and all three metrics on Pharos), DrugnomeAI retained a higher Spearman correlation on Recently Approved, and the other four comparisons were inconclusive. Since the pathway features do not record drug interactions, they do not qualify by the definition of label leakage we use here, so the main analysis retains them. Removing them as well lowered the precision@20 of DrugnomeAI further on Recently Approved and Pharos, to 0.82 and 0.31, and Varformer then exceeded DrugnomeAI in seven of the nine comparisons.

The gene inputs of Varformer contain no record of drug–gene interactions and no clinical precedence assessment. They do include four binary tractability indicators from Open Targets: a high-quality ligand in ChEMBL, a structure co-crystallized with a small molecule in the Protein Data Bank, a predicted high-quality pocket and a high-quality chemical probe (Methods). These indicators record whether chemical matter or a suitable binding site exists for the gene product. They do not meet our definition of leakage, because such properties are a precondition of small-molecule drug development and are present for many genes without a drug. Of the benchmark negatives, 15% have a high-quality ligand and 18% a co-crystal structure, compared with just 8% that have a DGIdb interaction.

## Supplementary Figures

**Supplementary Fig. 1:**
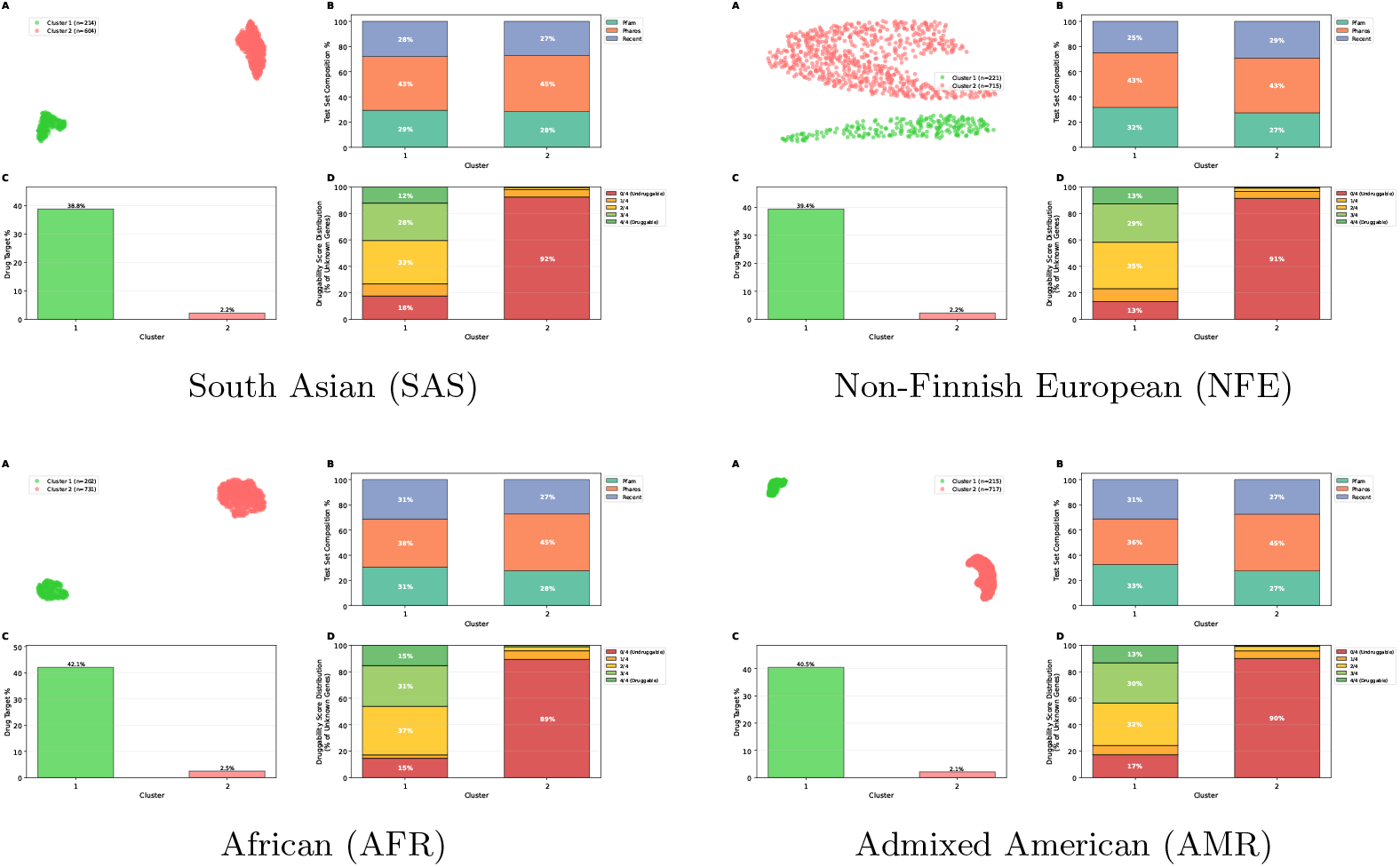
Druggability profiles across populations. Systematic evaluation of Varformer gene embeddings across the four ancestral cohorts. For each population, the four-panel figure displays: **A)** UMAP projection of the 256-dimensional latent space colored by unsupervised clusters (*k* = 2), **B)** the composition of benchmark test sets across clusters, **C)** the relative enrichment of drug targets, and **D)** the distribution of an independent gene-level druggability score (0–4) for unlabeled genes. Cluster 1 (green) is consistently enriched for tractable targets across all populations, while Cluster 2 (red) is dominated by “undruggable” genes (score 0/4), demonstrating that Varformer autonomously learns biologically and pharmacologically relevant representations.

**Supplementary Fig. 2:**
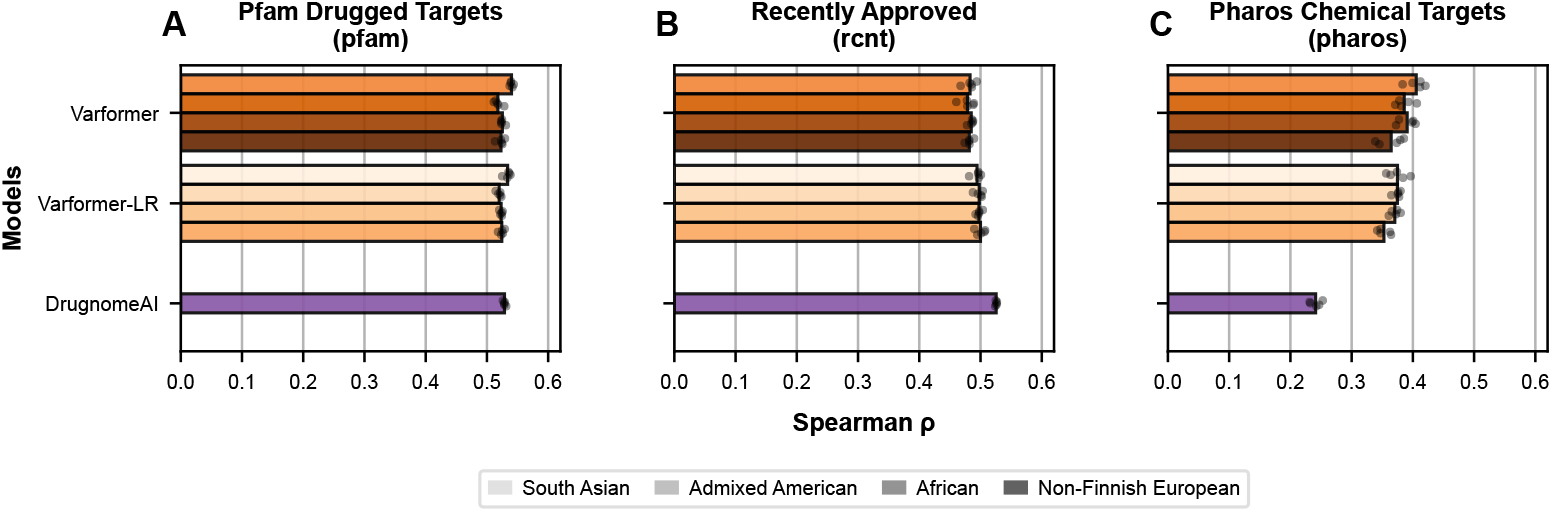
Spearman correlation on the three benchmarks by population. **A-C)** Spearman correlation between predicted scores and benchmark labels for Varformer, Varformer-LR and DrugnomeAI on the Pfam, Recently Approved and Pharos benchmarks. Bars show the mean across five seeds and points show individual seeds. Varformer and Varformer-LR are shown for each population; for DrugnomeAI, trained without drug–gene interaction features, each point is the mean of one seed across the four population-specific benchmark sets. Each value is computed once per run over the complete benchmark set.

**Supplementary Fig. 3:**
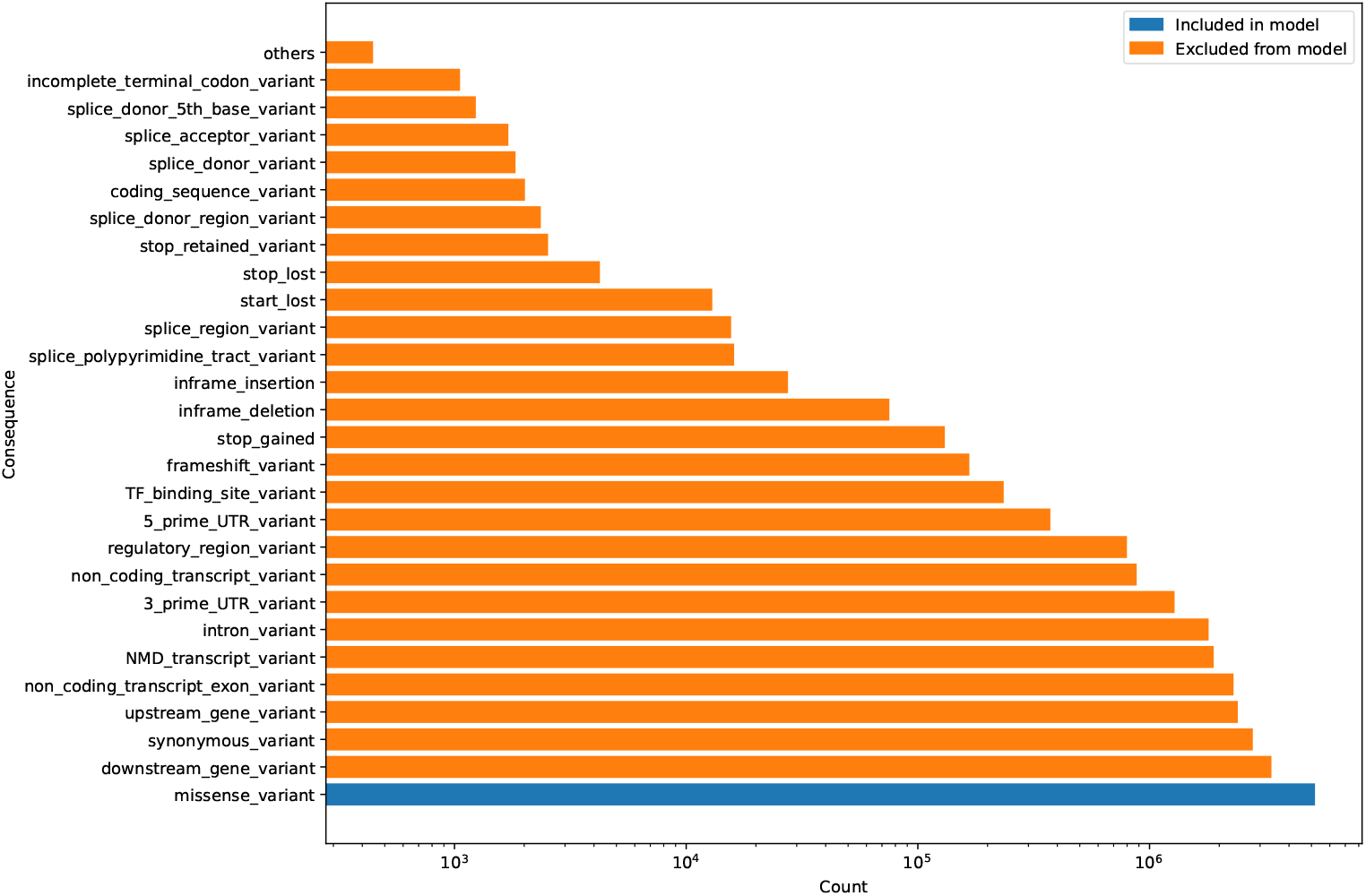
The counts of the potential variant consequences considered in our model. The ‘others’ column contains consequences that occur less than 1,000 times in the G&H dataset. These are: transcription factor binding site ablation variants (348), mature miRNA variants (40), protein altering variants (30), and start retained variants (7).

**Supplementary Fig. 4:**
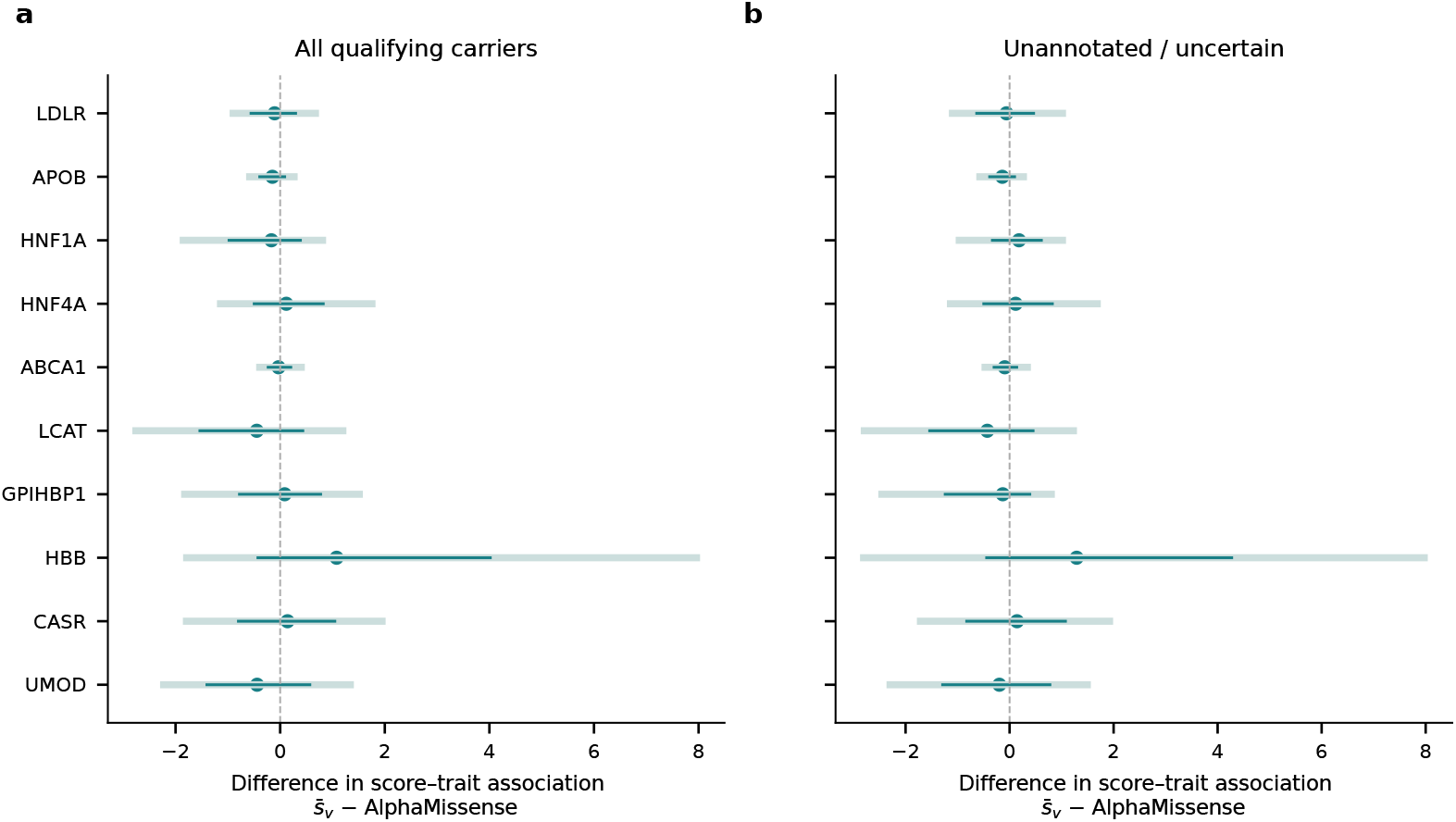
Exploratory paired phenotype comparisons. Gene-specific score–trait association differences between 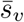 and AlphaMissense for **a)** all qualifying carriers and **b)** unannotated, uncertain or conflicting variants. Dark intervals are pointwise 95% variant-bootstrap intervals; pale intervals are approximate Bonferroni-adjusted intervals over 60 comparisons. All ten eligible genes are shown. No gene establishes superiority. Bootstrap draws: 50,000 per subset; carriers of each resampled variant remain together.

## Supplementary Tables

**Supplementary Table 1:** Performance metrics across test sets. Values are mean ± SD over 20 runs (5 seeds × 4 ancestral cohorts); all models are evaluated on the same seeds and therefore the same train/validation splits. Each metric is computed once per run over the complete benchmark set. The Mean ± SD column reports the spread across the three test-set means rather than across runs. Precision@20 uses a fixed cut of *k* = 20. The F1 score is computed with the *n*_+_ highest-scoring genes called positive, where *n*_+_ is the number of positives in the benchmark set. Tied scores are ordered by gene identifier. The random-baseline values are the analytic expectations for an arbitrary ranking: the positive rate for precision@20 and the F1 score, and zero for the Spearman correlation. DrugnomeAI is trained without drug–gene interaction features (Supplementary Note 9). The highest unrounded mean among the three scoring models is bolded.

| Model | Pfam | Recent | Pharos | Mean ± SD |
| --- | --- | --- | --- | --- |
| **Precision@20** |  |  |  |  |
| Varformer | **0.95 ± 0.04** | **0.86 ± 0.05** | **0.48 ± 0.08** | **0.76 ± 0.25** |
| Varformer-LR | **0.95 ± 0.05** | 0.80 ± 0.05 | 0.40 ± 0.07 | 0.72 ± 0.28 |
| DrugnomeAI | 0.91 ± 0.03 | 0.84 ± 0.04 | 0.39 ± 0.06 | 0.71 ± 0.28 |
| Random Baseline | 0.11 ± 0.00 | 0.12 ± 0.00 | 0.11 ± 0.00 | 0.11 ± 0.00 |
| **Spearman $\rho$** |  |  |  |  |
| Varformer | 0.53 ± 0.01 | 0.48 ± 0.01 | **0.39 ± 0.02** | **0.47 ± 0.07** |
| Varformer-LR | 0.53 ± 0.01 | 0.50 ± 0.01 | 0.37 ± 0.01 | 0.46 ± 0.08 |
| DrugnomeAI | **0.53 ± 0.01** | **0.53 ± 0.01** | 0.24 ± 0.01 | 0.43 ± 0.17 |
| Random Baseline | 0.00 | 0.00 | 0.00 | 0.00 |
| **F1 score** |  |  |  |  |
| Varformer | **0.87 ± 0.04** | **0.75 ± 0.04** | **0.44 ± 0.05** | **0.68 ± 0.22** |
| Varformer-LR | 0.83 ± 0.03 | 0.71 ± 0.04 | 0.39 ± 0.04 | 0.64 ± 0.23 |
| DrugnomeAI | 0.85 ± 0.03 | 0.74 ± 0.04 | 0.33 ± 0.04 | 0.64 ± 0.27 |
| Random Baseline | 0.11 ± 0.00 | 0.12 ± 0.00 | 0.11 ± 0.00 | 0.11 ± 0.00 |

**Supplementary Table 2:**
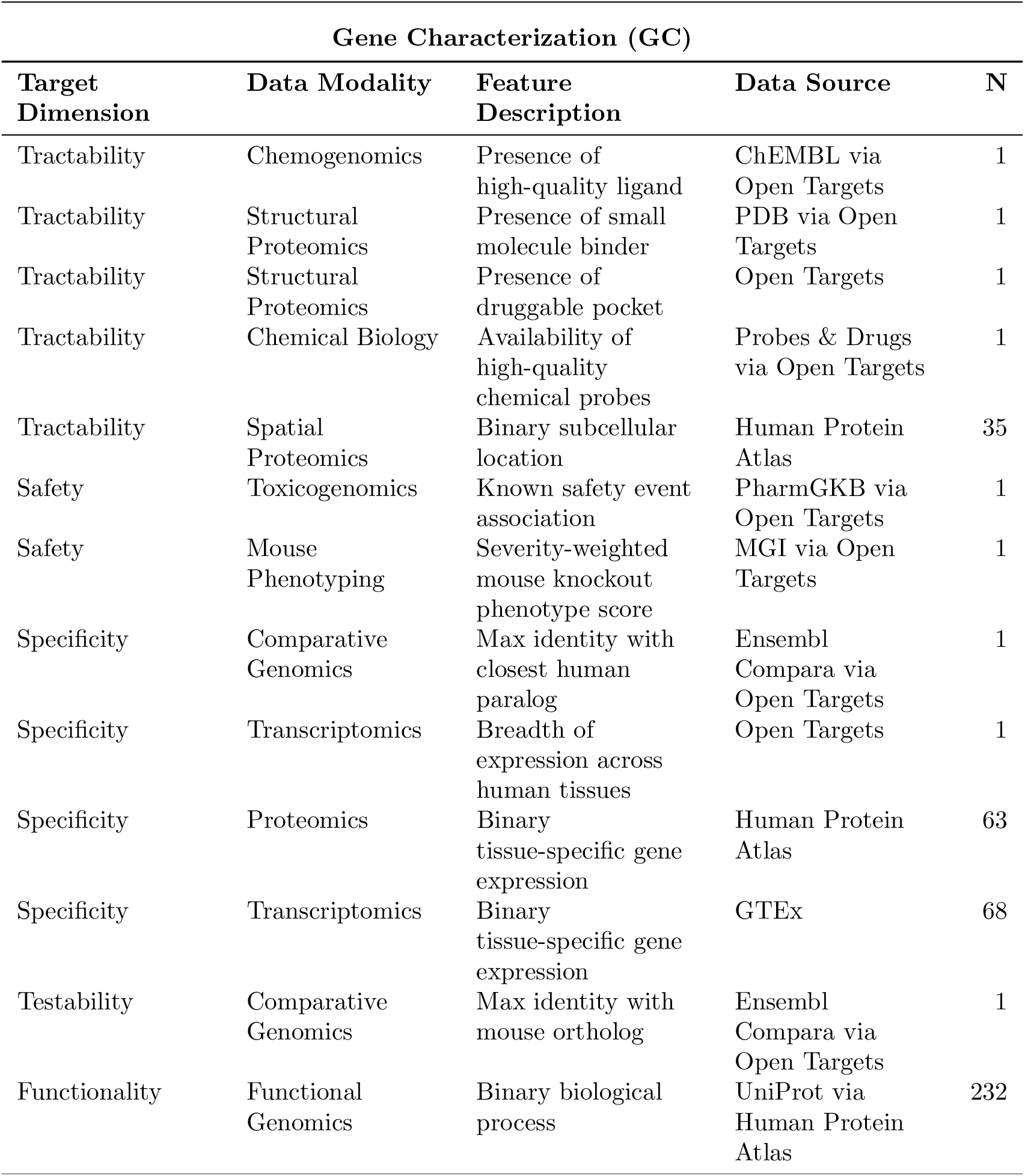
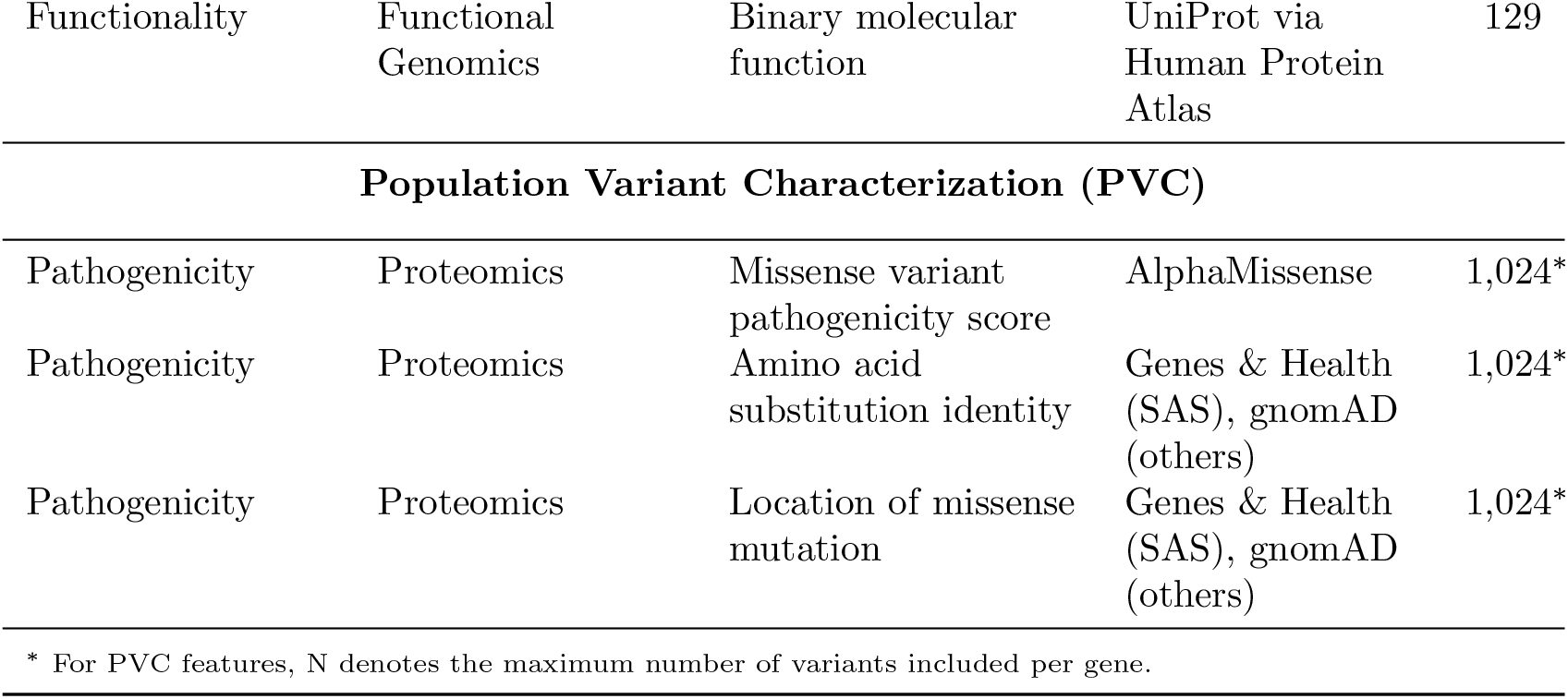
Description of the features used across Varformer’s different modules.

| Gene Characterization (GC) |  |  |  |  |
| --- | --- | --- | --- | --- |
| Target Dimension | Data Modality | Feature Description | Data Source | N |
| Tractability | Chemogenomics | Presence of high-quality ligand | ChEMBL via Open Targets | 1 |
| Tractability | Structural Proteomics | Presence of small molecule binder | PDB via Open Targets | 1 |
| Tractability | Structural Proteomics | Presence of druggable pocket | Open Targets | 1 |
| Tractability | Chemical Biology | Availability of high-quality chemical probes | Probes & Drugs via Open Targets | 1 |
| Tractability | Spatial Proteomics | Binary subcellular location | Human Protein Atlas | 35 |
| Safety | Toxicogenomics | Known safety event association | PharmGKB via Open Targets | 1 |
| Safety | Mouse Phenotyping | Severity-weighted mouse knockout phenotype score | MGI via Open Targets | 1 |
| Specificity | Comparative Genomics | Max identity with closest human paralog | Ensembl Compara via Open Targets | 1 |
| Specificity | Transcriptomics | Breadth of expression across human tissues | Open Targets | 1 |
| Specificity | Proteomics | Binary tissue-specific gene expression | Human Protein Atlas | 63 |
| Specificity | Transcriptomics | Binary tissue-specific gene expression | GTEX | 68 |
| Testability | Comparative Genomics | Max identity with mouse ortholog | Ensembl Compara via Open Targets | 1 |
| Functionality | Functional Genomics | Binary biological process | UniProt via Human Protein Atlas | 232 |
| Functionality | Functional Genomics | Binary molecular function | UniProt via Human Protein Atlas | 129 |
| **Population Variant Characterization (PVC)** |  |  |  |  |
| Pathogenicity | Proteomics | Missense variant pathogenicity score | AlphaMissense | 1,024* |
| Pathogenicity | Proteomics | Amino acid substitution identity | Genes & Health (SAS), gnomAD (others) | 1,024* |
| Pathogenicity | Proteomics | Location of missense mutation | Genes & Health (SAS), gnomAD (others) | 1,024* |
| * For PVC features, N denotes the maximum number of variants included per gene. |  |  |  |  |

**Supplementary Table 3:**
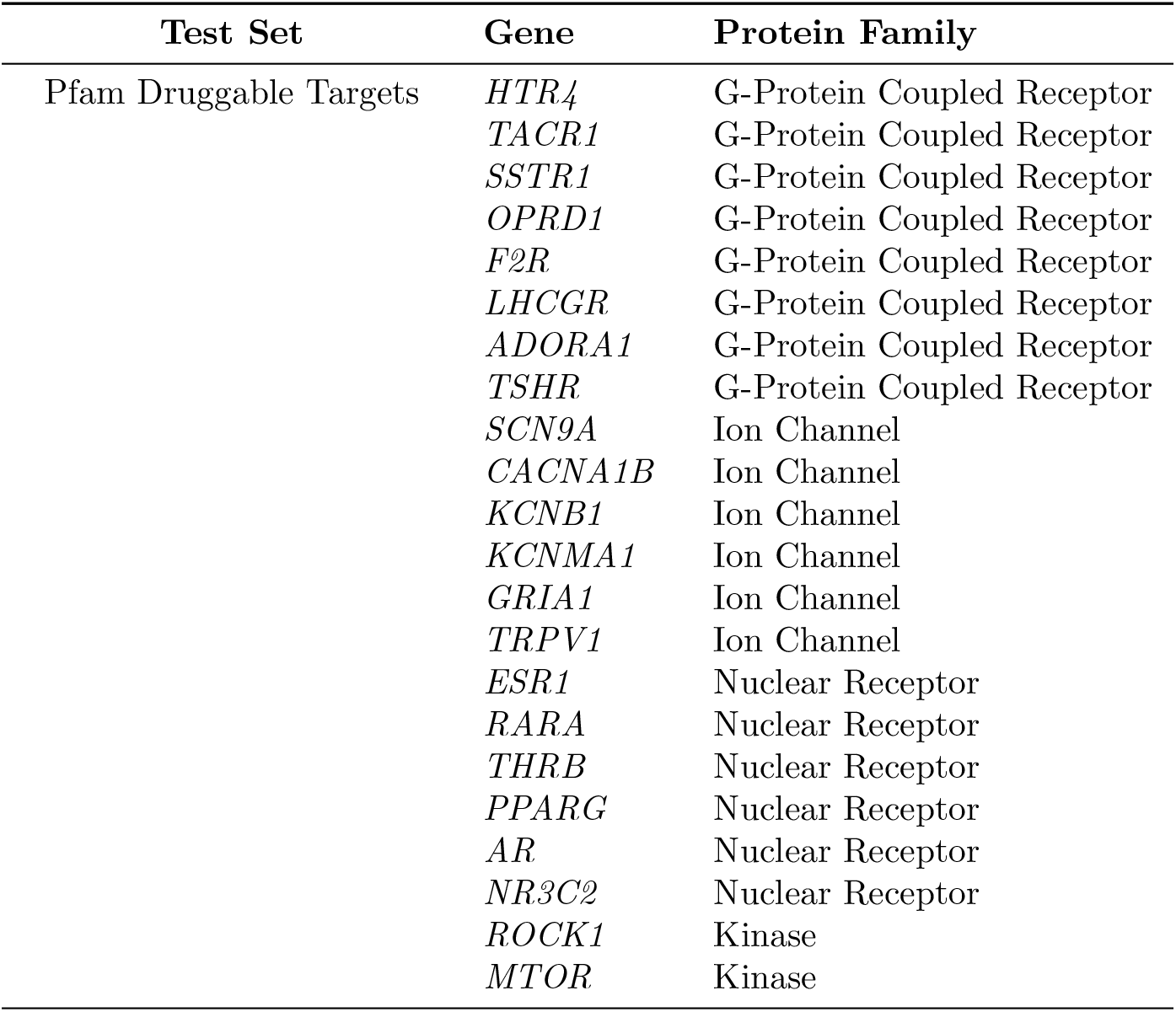
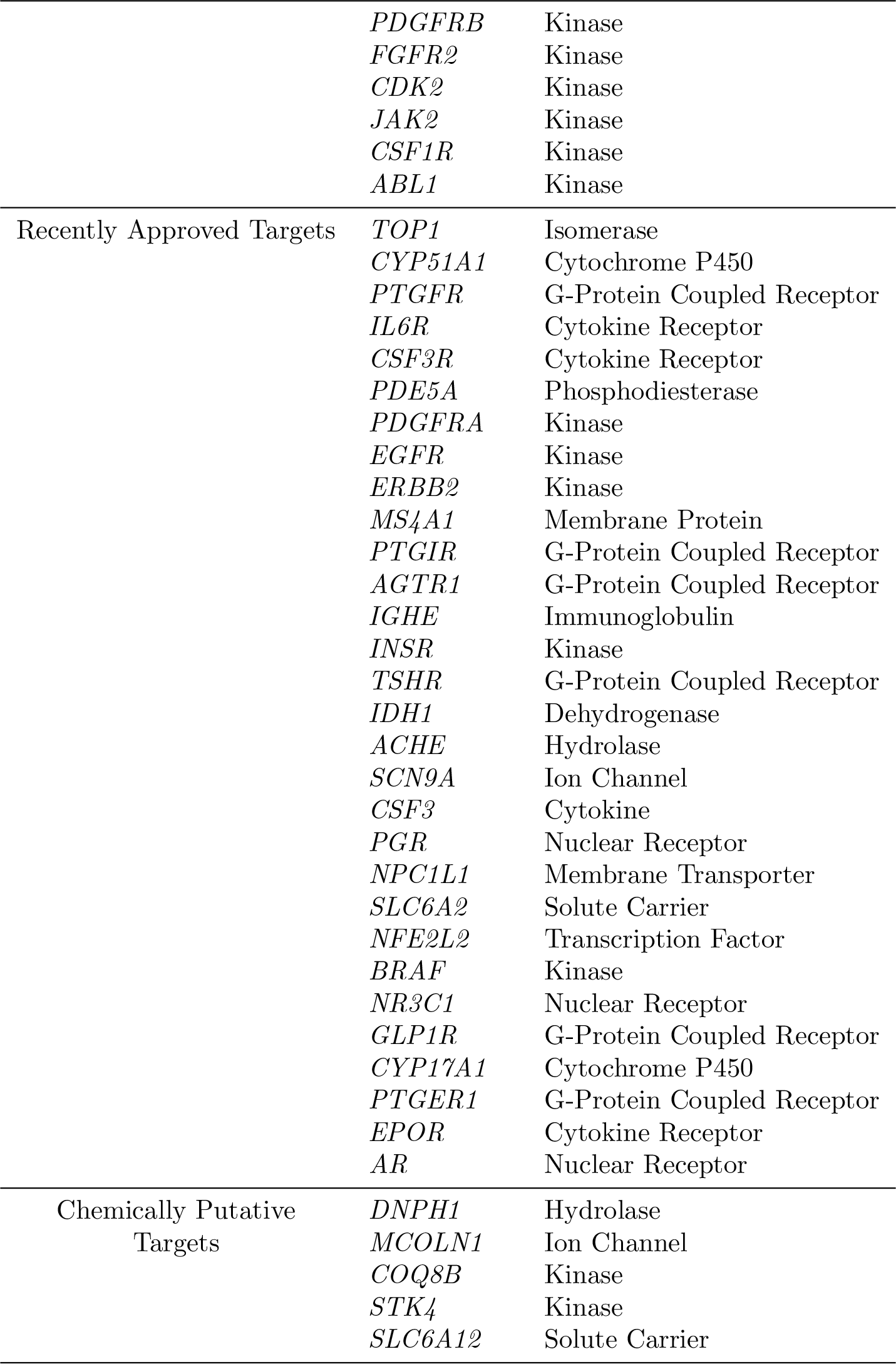
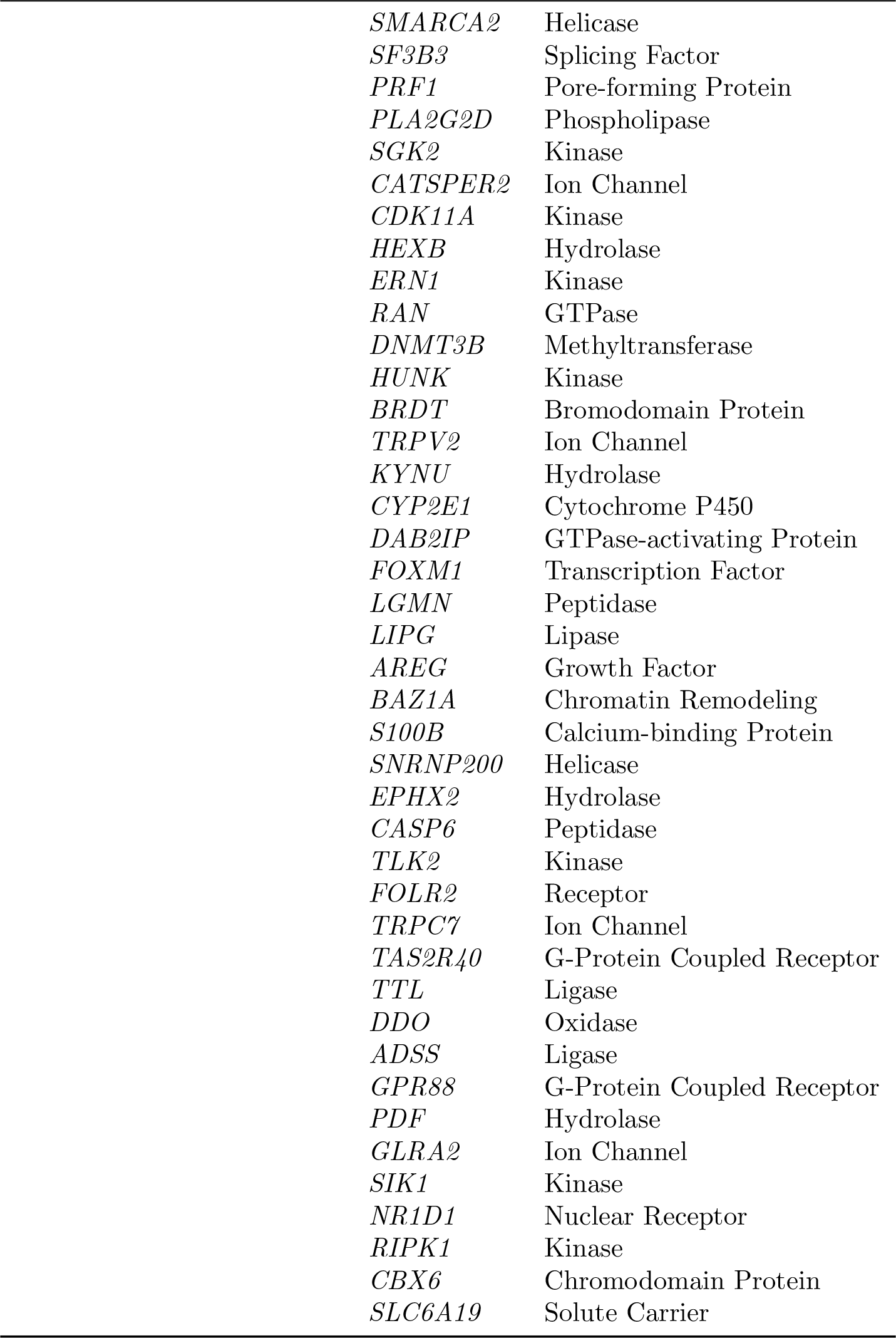
Test datasets comprising Pfam druggable targets, recently approved drug targets, and chemically putative targets with their respective protein families.

| Test Set | Gene | Protein Family |
| --- | --- | --- |
| Pfam Druggable Targets | *HTR4* | G-Protein Coupled Receptor |
|  | *TACR1* | G-Protein Coupled Receptor |
|  | *SSTR1* | G-Protein Coupled Receptor |
|  | *OPRD1* | G-Protein Coupled Receptor |
|  | *F2R* | G-Protein Coupled Receptor |
|  | *LHCGR* | G-Protein Coupled Receptor |
|  | *ADORA1* | G-Protein Coupled Receptor |
|  | *TSHR* | G-Protein Coupled Receptor |
|  | *SCN9A* | Ion Channel |
|  | *CACNA1B* | Ion Channel |
|  | *KCNB1* | Ion Channel |
|  | *KCNMA1* | Ion Channel |
|  | *GRIA1* | Ion Channel |
|  | *TRPV1* | Ion Channel |
|  | *ESR1* | Nuclear Receptor |
|  | *RARA* | Nuclear Receptor |
|  | *THRB* | Nuclear Receptor |
|  | *PPARG* | Nuclear Receptor |
|  | *AR* | Nuclear Receptor |
|  | *NR3C2* | Nuclear Receptor |
|  | *ROCK1* | Kinase |
|  | *MTOR* | Kinase |

| **Test Set** | **Gene** | **Protein Family** |
| --- | --- | --- |
|  | *PDGFRB* | Kinase |
|  | *FGFR2* | Kinase |
|  | *CDK2* | Kinase |
|  | *JAK2* | Kinase |
|  | *CSF1R* | Kinase |
|  | *ABL1* | Kinase |
| Recently Approved Targets | *TOP1* | Isomerase |
|  | *CYP51A1* | Cytochrome P450 |
|  | *PTGFR* | G-Protein Coupled Receptor |
|  | *IL6R* | Cytokine Receptor |
|  | *CSF3R* | Cytokine Receptor |
|  | *PDE5A* | Phosphodiesterase |
|  | *PDGFRA* | Kinase |
|  | *EGFR* | Kinase |
|  | *ERBB2* | Kinase |
|  | *MS4A1* | Membrane Protein |
|  | *PTGIR* | G-Protein Coupled Receptor |
|  | *AGTR1* | G-Protein Coupled Receptor |
|  | *IGHE* | Immunoglobulin |
|  | *INSR* | Kinase |
|  | *TSHR* | G-Protein Coupled Receptor |
|  | *IDH1* | Dehydrogenase |
|  | *ACHE* | Hydrolase |
|  | *SCN9A* | Ion Channel |
|  | *CSF3* | Cytokine |
|  | *PGR* | Nuclear Receptor |
|  | *NPC1L1* | Membrane Transporter |
|  | *SLC6A2* | Solute Carrier |
|  | *NFE2L2* | Transcription Factor |
|  | *BRAF* | Kinase |
|  | *NR3C1* | Nuclear Receptor |
|  | *GLP1R* | G-Protein Coupled Receptor |
|  | *CYP17A1* | Cytochrome P450 |
|  | *PTGER1* | G-Protein Coupled Receptor |
|  | *EPOR* | Cytokine Receptor |
|  | *AR* | Nuclear Receptor |
| Chemically Putative Targets | *DNPH1* | Hydrolase |
|  | *MCOLN1* | Ion Channel |
|  | *COQ8B* | Kinase |
|  | *STK4* | Kinase |
|  | *SLC6A12* | Solute Carrier |
|  | *SMARCA2* | Helicase |
|  | *SF3B3* | Splicing Factor |
|  | *PRF1* | Pore-forming Protein |
|  | *PLA2G2D* | Phospholipase |
|  | *SGK2* | Kinase |
|  | *CATSPER2* | Ion Channel |
|  | *CDK11A* | Kinase |
|  | *HEXB* | Hydrolase |
|  | *ERN1* | Kinase |
|  | *RAN* | GTPase |
|  | *DNMT3B* | Methyltransferase |
|  | *HUNK* | Kinase |
|  | *BRDT* | Bromodomain Protein |
|  | *TRPV2* | Ion Channel |
|  | *KYNU* | Hydrolase |
|  | *CYP2E1* | Cytochrome P450 |
|  | *DAB2IP* | GTPase-activating Protein |
|  | *FOXM1* | Transcription Factor |
|  | *LGMN* | Peptidase |
|  | *LIPG* | Lipase |
|  | *AREG* | Growth Factor |
|  | *BAZ1A* | Chromatin Remodeling |
|  | *S100B* | Calcium-binding Protein |
|  | *SNRNP200* | Helicase |
|  | *EPHX2* | Hydrolase |
|  | *CASP6* | Peptidase |
|  | *TLK2* | Kinase |
|  | *FOLR2* | Receptor |
|  | *TRPC7* | Ion Channel |
|  | *TAS2R40* | G-Protein Coupled Receptor |
|  | *TTL* | Ligase |
|  | *DDO* | Oxidase |
|  | *ADSS* | Ligase |
|  | *GPR88* | G-Protein Coupled Receptor |
|  | *PDF* | Hydrolase |
|  | *GLRA2* | Ion Channel |
|  | *SIK1* | Kinase |
|  | *NR1D1* | Nuclear Receptor |
|  | *RIPK1* | Kinase |
|  | *CBX6* | Chromodomain Protein |
|  | *SLC6A19* | Solute Carrier |

**Supplementary Table 4:**
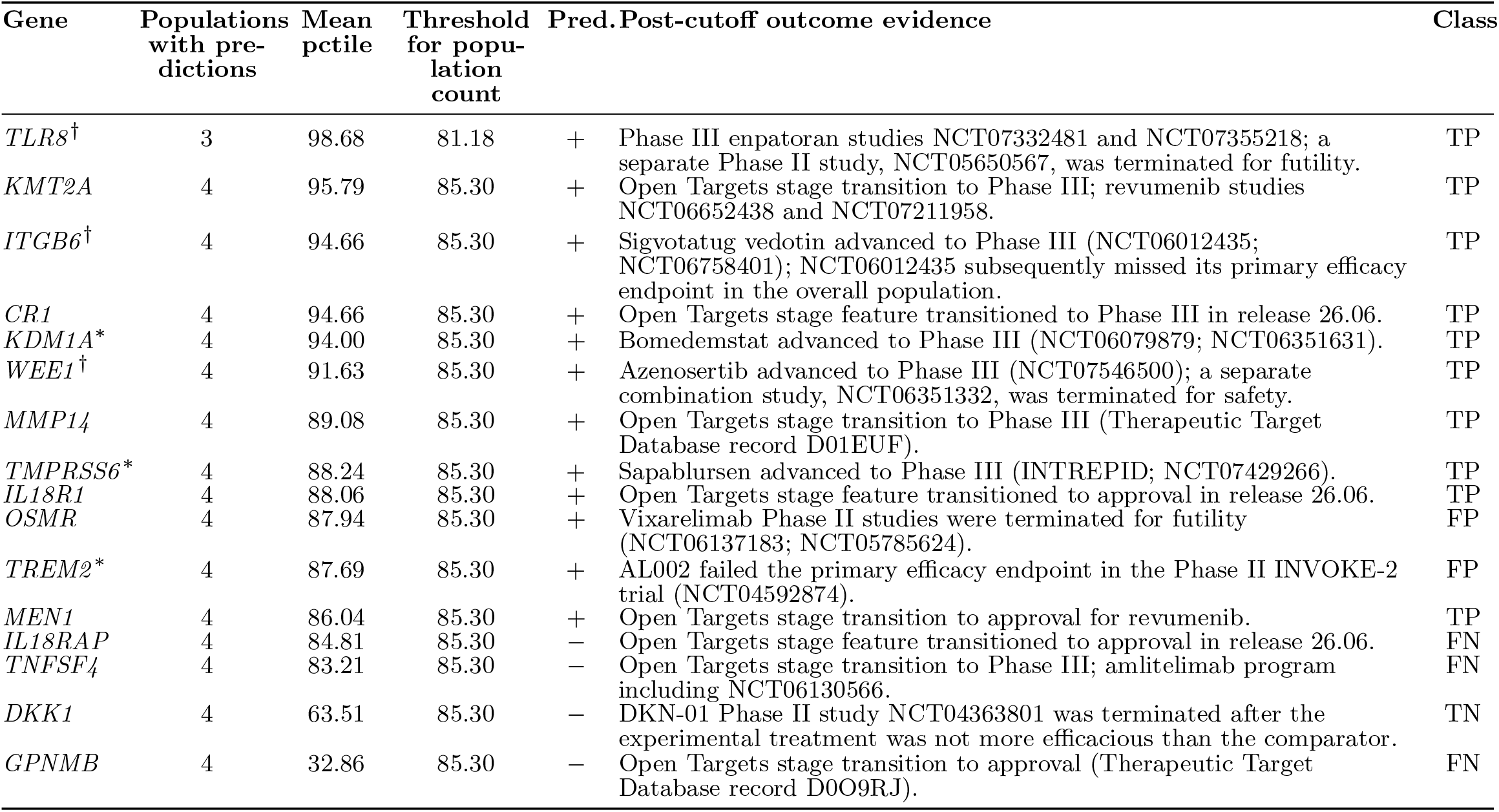
Prospective outcome audit using cross-population Varformer scores. Genes are ordered by mean cross-population percentile. The table reports the number of populations with an available five-seed ensemble prediction for each gene. The threshold for each reported population count is the 91.1th percentile estimated among training-unlabeled genes with predictions in that number of populations: 48.50 for two populations, 81.18 for three populations, and 85.30 for all four populations. The positive outcome is an Open Targets target-stage transition from Phase I/II in release 23.06 to Phase III or approval in release 26.06; the negative outcome requires a manually verified, directly target-attributable efficacy or safety failure. Unresolved programs are excluded. Eleven *IFNA* mappings to one sifalimumab program are excluded from this primary table. The unit is a target mapping rather than an independent program. ^*^ Main-text case study (§6); ^†^ the target also had a separate efficacy or safety setback, illustrating a mixed program history. Class: TP, stage progression with a score above the threshold for the listed number of populations; TN, confirmed failure with a score below that threshold; FP, confirmed failure with a score above that threshold; FN, stage progression despite a score below that threshold. *TREM2* has no clinical phase annotation in release 23.06 and was included on the basis of its registered Phase II trial.

**Supplementary Table 5:** Exploratory retrieval of known pathogenic variants on 1,164 carrier–variant pairs. Known pathogenic variants include ClinVar pathogenic and likely pathogenic classifications. Entries are recall percentages within the scoreable subset, not whole-exome sensitivity. AM denotes AlphaMissense; AF in combinations denotes negated frequency. Combination weights are equal.

| Score | @1 | @5 | @10 | @20 | @50 | @100 |
| --- | --- | --- | --- | --- | --- | --- |
| $\bar{s}_v$ | 6.3 | 23.4 | 36.2 | 56.7 | 81.7 | 94.5 |
| AlphaMissense | 5.0 | 21.3 | 36.2 | 53.3 | 80.0 | 95.7 |
| CADD | 2.1 | 15.9 | 30.6 | 51.5 | 84.4 | 97.2 |
| REVEL | 17.3 | 46.8 | 59.5 | 71.5 | 82.3 | 89.3 |
| Negated AF | 0.0 | 1.7 | 4.5 | 12.5 | 38.2 | 73.5 |
| Gene score | 0.6 | 7.2 | 16.4 | 34.0 | 66.1 | 88.1 |
| $\bar{s}_v$+AF | 3.4 | 11.9 | 23.0 | 36.3 | 67.4 | 92.2 |
| AM+AF | 2.1 | 11.9 | 21.7 | 36.3 | 67.2 | 92.3 |
| AM+AF+$\bar{s}_v$ | 3.4 | 14.6 | 27.5 | 47.3 | 79.2 | 94.0 |

**Supplementary Table 6:**
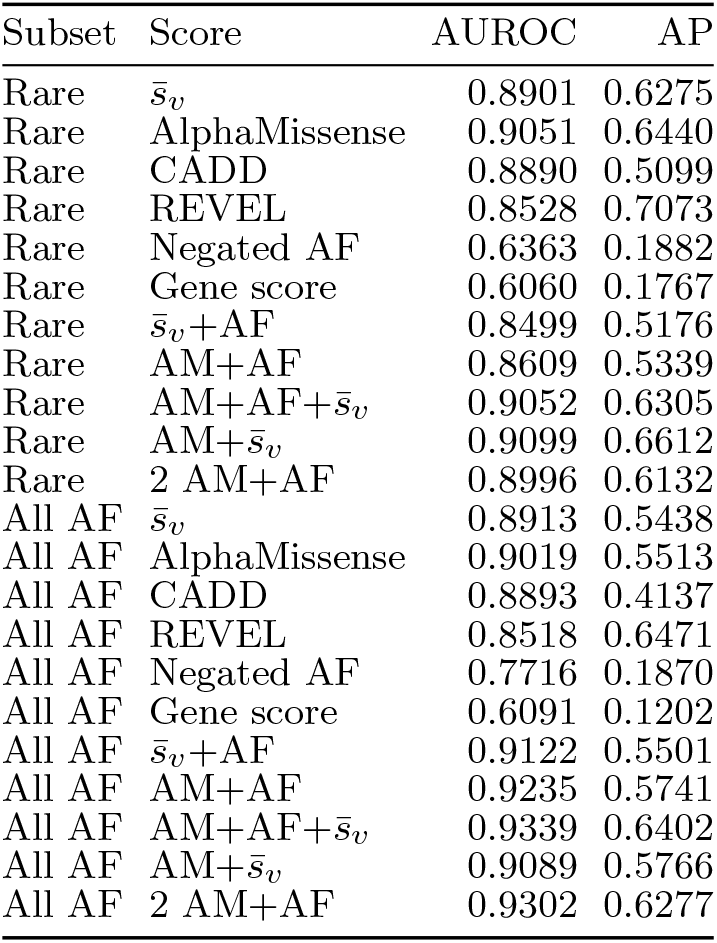
Known-label discrimination among scoreable G&H DNA variants. Rare: AF *<* 10^−3^, 2,477 pathogenic and 15,580 benign variants. All AF: 2,521 pathogenic and 25,777 benign variants. AP is average precision. AM denotes AlphaMissense; AF denotes allele rarity. Combinations use equal percentile weights, except 2 AM+AF, which assigns two-thirds weight to AM and one-third to AF. Unannotated, uncertain and conflicting variants are excluded. These labels overlap the evidence used in retrieval.

**Supplementary Table 7:**
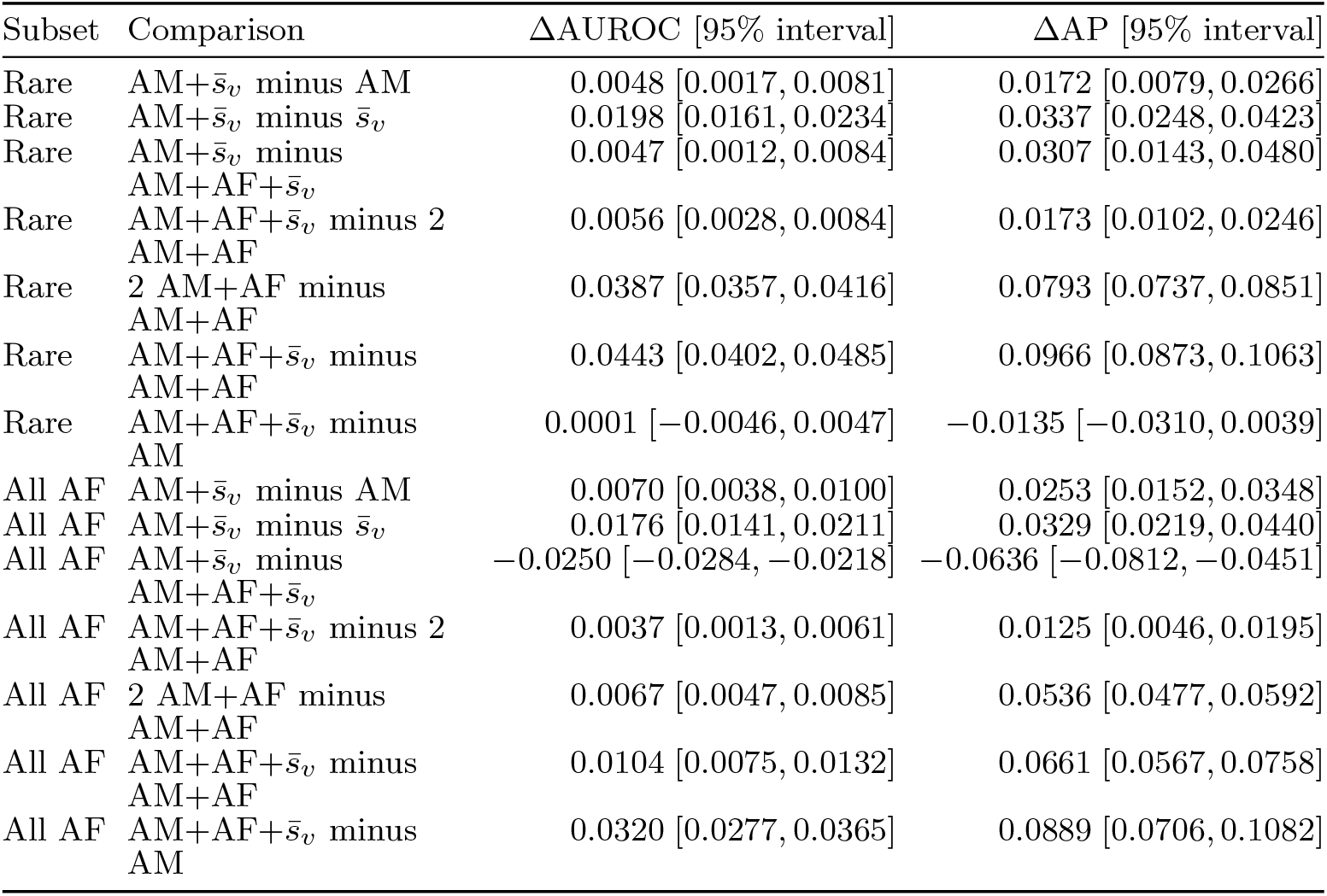
Paired classification differences for G&H score combinations. Entries are the observed difference followed by its pointwise 95% interval from 1,000 paired gene-bootstrap draws. Positive values favor the first method. AM denotes AlphaMissense; AF denotes allele rarity. Equal-weight percentile combinations are compared with a control, 2 AM+AF, assigning two-thirds weight to AM and one-third to AF. These exploratory intervals are not adjusted for multiple comparisons; the rare and all-frequency sets overlap.

| Subset | Comparison | ΔAUROC [95% interval] | ΔAP [95% interval] |
| --- | --- | --- | --- |
| Rare | AM+$\bar{s}_v$ minus AM | 0.0048 [0.0017, 0.0081] | 0.0172 [0.0079, 0.0266] |
| Rare | AM+$\bar{s}_v$ minus $\bar{s}_v$ | 0.0198 [0.0161, 0.0234] | 0.0337 [0.0248, 0.0423] |
| Rare | AM+$\bar{s}_v$ minus<br>AM+AF+$\bar{s}_v$ | 0.0047 [0.0012, 0.0084] | 0.0307 [0.0143, 0.0480] |
| Rare | AM+AF+$\bar{s}_v$ minus 2<br>AM+AF | 0.0056 [0.0028, 0.0084] | 0.0173 [0.0102, 0.0246] |
| Rare | 2 AM+AF minus<br>AM+AF | 0.0387 [0.0357, 0.0416] | 0.0793 [0.0737, 0.0851] |
| Rare | AM+AF+$\bar{s}_v$ minus<br>AM+AF | 0.0443 [0.0402, 0.0485] | 0.0966 [0.0873, 0.1063] |
| Rare | AM+AF+$\bar{s}_v$ minus<br>AM | 0.0001 [−0.0046, 0.0047] | −0.0135 [−0.0310, 0.0039] |
| All AF | AM+$\bar{s}_v$ minus AM | 0.0070 [0.0038, 0.0100] | 0.0253 [0.0152, 0.0348] |
| All AF | AM+$\bar{s}_v$ minus $\bar{s}_v$ | 0.0176 [0.0141, 0.0211] | 0.0329 [0.0219, 0.0440] |
| All AF | AM+$\bar{s}_v$ minus<br>AM+AF+$\bar{s}_v$ | −0.0250 [−0.0284, −0.0218] | −0.0636 [−0.0812, −0.0451] |
| All AF | AM+AF+$\bar{s}_v$ minus 2<br>AM+AF | 0.0037 [0.0013, 0.0061] | 0.0125 [0.0046, 0.0195] |
| All AF | 2 AM+AF minus<br>AM+AF | 0.0067 [0.0047, 0.0085] | 0.0536 [0.0477, 0.0592] |
| All AF | AM+AF+$\bar{s}_v$ minus<br>AM+AF | 0.0104 [0.0075, 0.0132] | 0.0661 [0.0567, 0.0758] |
| All AF | AM+AF+$\bar{s}_v$ minus<br>AM | 0.0320 [0.0277, 0.0365] | 0.0889 [0.0706, 0.1082] |

**Supplementary Table 8:**
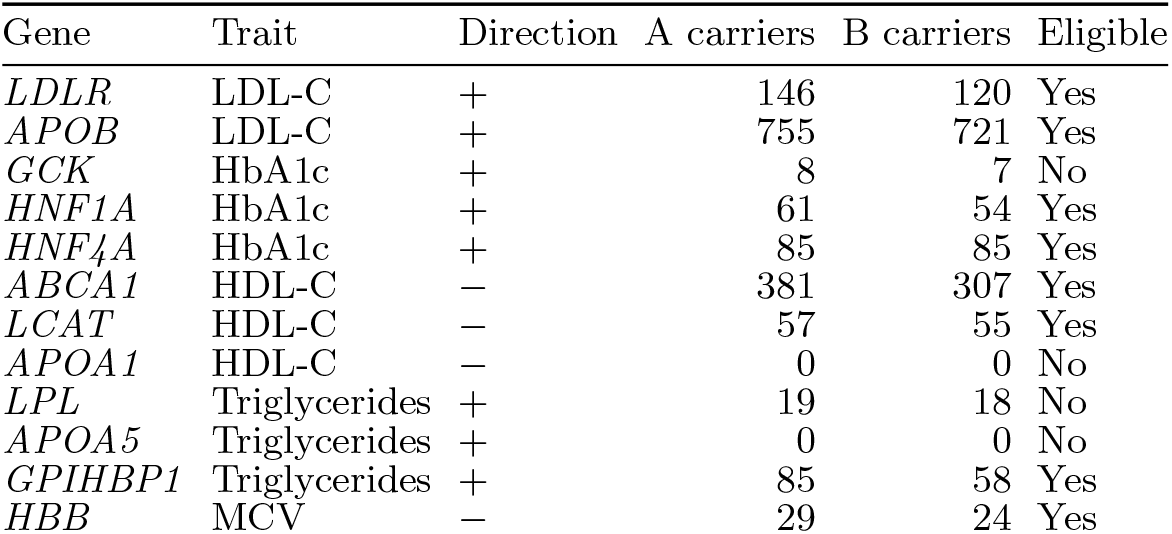
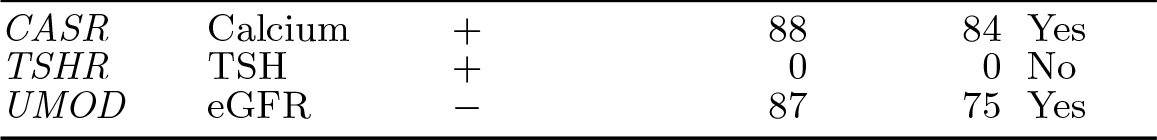
Gene–trait panel and observed carrier counts. Direction orients traits toward the hypothesized effect of damaging variation. A: all qualifying carriers; B: unannotated, uncertain or conflicting variants. The final column denotes eligibility for the exploratory gene-specific follow-up in both subsets. Counts of 1–5 remain suppressed. APOB, CASR and TSHR can have bidirectional variant effects.

**Supplementary Table 9:** Pooled score–trait associations with gene fixed effects. A: 1,801 carrier observations and 723 variants; B: 1,608 observations and 675 variants. Intervals are pointwise 95% within-gene variant-bootstrap intervals. Combination results are secondary. Intervals for the pooled 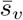 association include zero in both subsets.

| Subset | Score | Score–trait association | 95% interval |
| --- | --- | --- | --- |
| A | $\bar{s}_v$ | 0.196 | [−0.009, 0.402] |
| A | AlphaMissense | 0.294 | [0.087, 0.501] |
| A | CADD | 0.240 | [0.038, 0.435] |
| A | REVEL | 0.194 | [−0.025, 0.403] |
| A | Negated AF | 0.496 | [0.239, 0.761] |
| A | $\bar{s}_v$+AF | 0.407 | [0.154, 0.663] |
| A | AM+AF | 0.473 | [0.231, 0.721] |
| A | AM+AF+$\bar{s}_v$ | 0.338 | [0.121, 0.555] |
| B | $\bar{s}_v$ | 0.143 | [−0.074, 0.356] |
| B | AlphaMissense | 0.249 | [0.030, 0.463] |
| B | CADD | 0.194 | [−0.026, 0.403] |
| B | REVEL | 0.195 | [−0.035, 0.410] |
| B | Negated AF | 0.421 | [0.163, 0.679] |
| B | $\bar{s}_v$+AF | 0.329 | [0.058, 0.587] |
| B | AM+AF | 0.395 | [0.141, 0.643] |
| B | AM+AF+$\bar{s}_v$ | 0.272 | [0.039, 0.496] |

**Supplementary Table 10:**
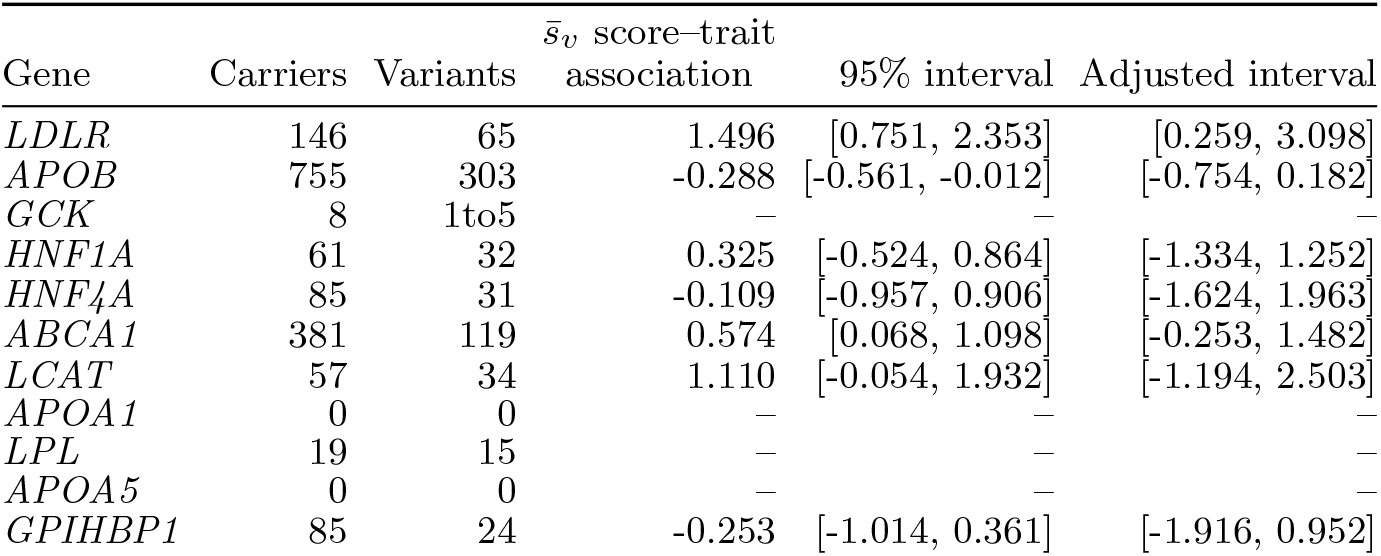
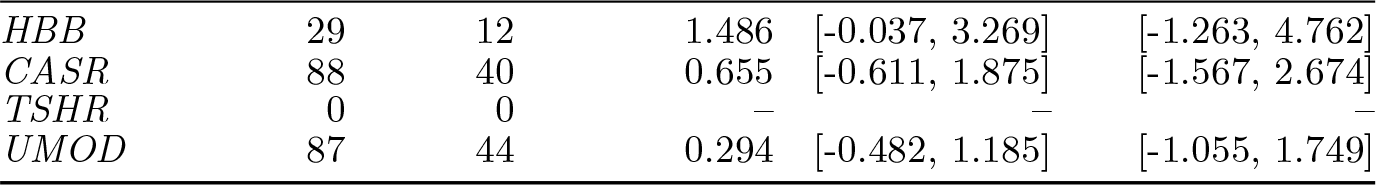
Exploratory gene-specific phenotype results, subset Carrier and distinct-variant counts precede the 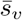 score–trait association and its intervals. The adjusted interval uses the 60-comparison family. Dashes indicate insufficient support; no missing estimate is treated as zero. Source: approved screenshot-transcribed aggregates.

**Supplementary Table 11:** Exploratory gene-specific phenotype results, subset Carrier and distinct-variant counts precede the 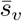 score–trait association and its intervals. The adjusted interval uses the 60-comparison family. Dashes indicate insufficient support; no missing estimate is treated as zero. Source: approved screenshot-transcribed aggregates.

| Gene | Carriers | Variants | $\bar{s}_v$ score-trait | | |
| --- | --- | --- | --- | --- | --- |
|  |  |  | association | 95% interval | Adjusted interval |
| *LDLR* | 120 | 56 | 1.437 | [0.583, 2.560] | [0.004, 3.639] |
| *APOB* | 721 | 289 | -0.299 | [-0.575, -0.023] | [-0.775, 0.170] |
| *GCK* | 7 | 1to5 | — | — | — |
| *HNF1A* | 54 | 28 | 0.326 | [-0.501, 0.907] | [-1.190, 1.316] |
| *HNF4A* | 85 | 31 | -0.109 | [-0.949, 0.903] | [-1.619, 1.817] |
| *ABCA1* | 307 | 110 | 0.522 | [-0.015, 1.053] | [-0.379, 1.434] |
| *LCAT* | 55 | 32 | 1.101 | [-0.071, 1.934] | [-1.207, 2.493] |
| *APOA1* | 0 | 0 | — | — | — |
| *LPL* | 18 | 14 | — | — | — |
| *APOA5* | 0 | 0 | — | — | — |
| *GPIHBP1* | 58 | 21 | -0.669 | [-1.601, 0.011] | [-2.615, 0.589] |
| *HBB* | 24 | 11 | 1.369 | [-0.305, 3.072] | [-1.982, 3.891] |
| *CASR* | 84 | 39 | 0.642 | [-0.643, 1.857] | [-1.591, 2.665] |
| *TSHR* | 0 | 0 | — | — | — |
| *UMOD* | 75 | 41 | 0.484 | [-0.430, 1.269] | [-1.161, 1.740] |

**Supplementary Table 12:**
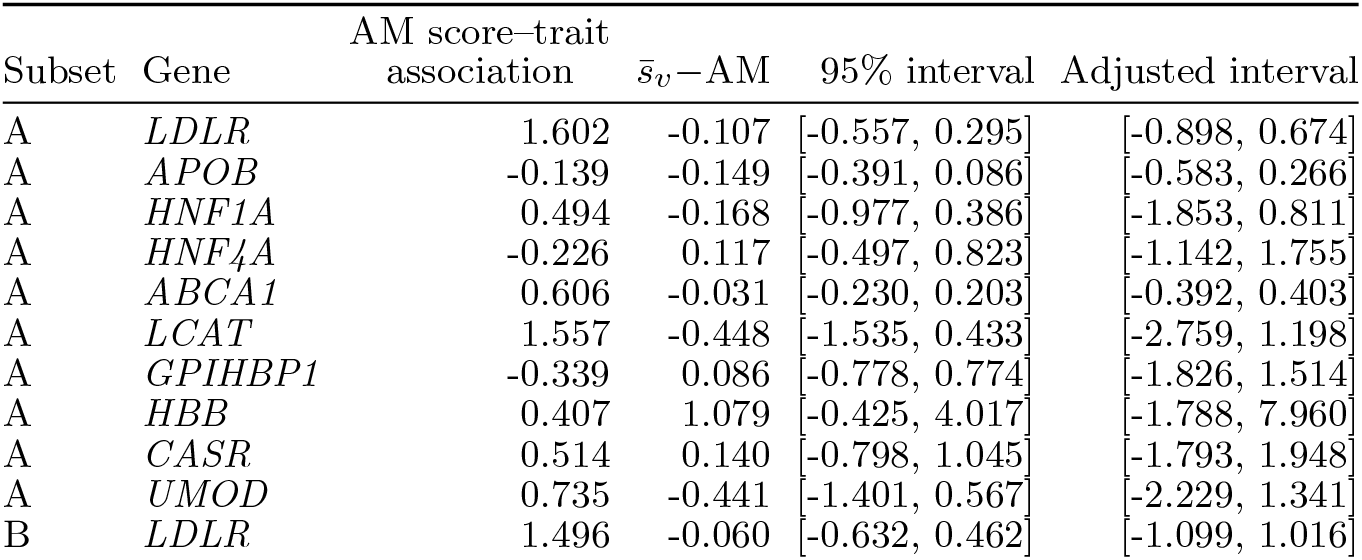
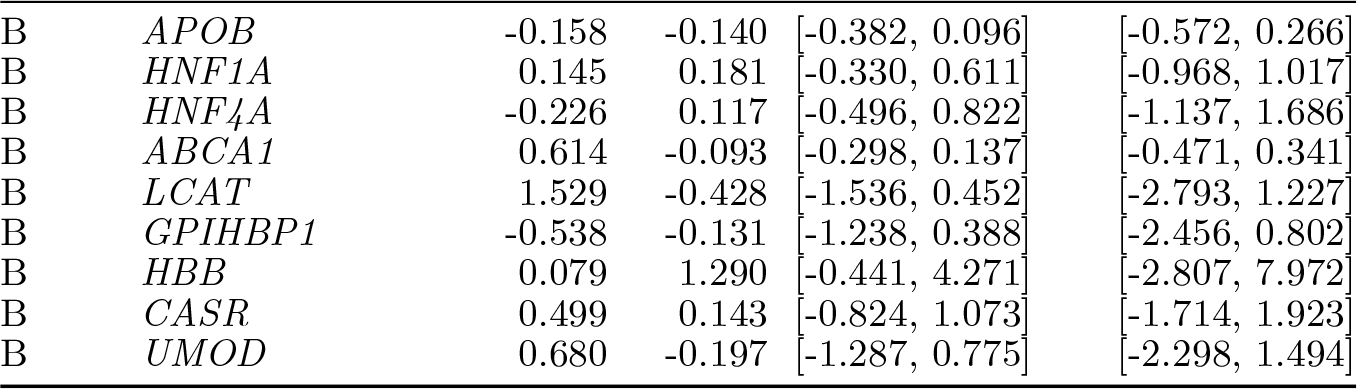
Exploratory paired phenotype comparisons. AM denotes AlphaMissense. Intervals concern the paired difference in score–trait association, not separate confidence intervals compared by eye. All 20 eligible gene/subset combinations are retained.

